# From literature to biodiversity data: mining arthropod organismal and ecological traits with machine learning

**DOI:** 10.1101/2025.02.18.638830

**Authors:** Joseph Cornelius, Harald Detering, Oscar Lithgow-Serrano, Donat Agosti, Fabio Rinaldi, Robert M. Waterhouse

## Abstract

The fields of taxonomy and biodiversity research have witnessed an exponential growth in published literature. This vast corpus of articles holds information on the diverse biological traits of organisms and their ecologies. However, access to and extraction of relevant data from this extensive resource remain challenging. Advances in text and data mining (TDM) and Natural Language Processing (NLP) techniques offer new opportunities for liberating such information from the literature. Testing and using such approaches to annotate articles in machine actionable formats is therefore necessary to enable the exploitation of existing knowledge in new biology, ecology, and evolution research. Here we explore the potential of these methods to annotate and extract organismal and ecological trait data for the most diverse animal group on Earth, the arthropods. The article processing workflow uses manually curated trait dictionaries with trained NLP models to perform labelling of entities and relationships of thousands of articles. A subset of manually annotated documents facilitated the formal evaluation of the performance of the workflow in terms of entity recognition and normalisation, and relationship extraction, highlighting several important technical challenges. The results are made available to the scientific community through an interactive web tool and queryable resource, the ArTraDB Arthropod Trait Database. These methodological explorations provide a framework that could be extended beyond the arthropods, where TDM and NLP approaches applied to the taxonomy and biodiversity literature will greatly facilitate data synthesis studies and literature reviews, the identification of knowledge gaps and biases, as well as the data-informed investigation of ecological and evolutionary trends and patterns.

## Introduction

The existing detailed knowledge on biodiversity and the natural world is contained largely in the form of an extensive and growing corpus of scientific publications (McCallen et al. 2019). This knowledge harbours important insights to better understand the dynamics and dimensions of major challenges facing our planet today, such as the global biodiversity crisis and the impact of climate change on the distribution of species. Large portions of that information have been difficult to access because they are unstructured, in printed formats, including portable data format (PDF), which have been difficult to machine operate, or which are behind paywalled access. With increasing digitisation of the scientific publishing process, and thanks to comprehensive digitisation efforts of natural history collections (Hedrick et al. 2020), an increasing number of documents have become digitally accessible. For example, PubMedCentral (PMC) alone contains millions of machine actionable articles (Rosonovski et al. 2023), including millions of supplementary data files, and tens of millions of abstracts are accessible through PubMed. These documents can be used for text and data mining (TDM) and natural language processing (NLP) to better annotate the literature, offering opportunities to liberate data and knowledge from publications. However, even though such literature mining approaches are recognised as important tools in biology, ecology, and evolution research, their potential is currently far from fully realised (Farrell et al. 2022, 2024).

To focus the methodological explorations of these opportunities, aiming to annotate and extract organismal and ecological trait data for the most numerous and diverse animal group on Earth with estimates of 6.8 million terrestrial species (Stork 2018), the phylum Arthropoda represents an excellent case study. Arthropods have fascinated researchers and amateur entomologists for centuries, leading to a vast accumulation of knowledge about how countless evolutionary adaptations have enabled them to exploit so many ecological niches (Grimaldi and Engel 2005). The most extensive knowledge is often biased towards the more charismatic species and those that serve as models in research, or that impact human health and agriculture. The decades of accrued learning also present substantial variability in terms of what types of trait data have been collected and with which methodologies (Wong et al. 2019). More recently, this biological knowledge is being extended through the acquisition of increasing amounts of genomics data to explore the genetics underlying these traits (Feron and Waterhouse 2022b). The motivation is to understand how genetic and genomic changes relate to observable phenotypic differences, *i.e.* traits, amongst species. Thanks to ongoing developments in bioinformatics, comparative genomics analyses are generally scalable to increasingly larger datasets, taking advantage of rapidly accumulating numbers of species with sequenced genomes (Feron and Waterhouse 2022a). However, this is not being matched by equivalent advances in the cataloguing of species traits for which manual collection and curation efforts cannot keep pace with the needs to access trait data for large-scale quantitative analyses. Literature mining offers potential solutions to overcoming these challenges, for example, building a database of insect egg size and shape for more than 6’700 species relied on information extracted from 1’756 publications (Church et al. 2019), cataloguing traits of 12’448 butterfly and moth species involved extracting information from 117 field guides and species accounts (Shirey et al. 2022), and compiling an expert-curated trait database of 520 subterranean spiders examined 255 taxonomic descriptions from the World Spider Catalog and the Spiders of Europe repository (Mammola et al. 2022). Therefore, efforts to develop systematic approaches for mining the literature to build comprehensive open databases of species’ organismal and ecological traits should increasingly provide researchers with direct access to much-needed large-scale biodiversity data.

Here, we present a TDM and NLP framework for the automated labelling of identified arthropod species (taxa), their organismal and ecological traits, and the associated trait values in taxonomic and biodiversity research articles. We focus on PMC articles containing taxonomic treatments of arthropods, *i.e.* structured sections of publications that describe and define the name and features of species, leveraging the large resource of Plazi’s TreatmentBank (Guidoti et al. 2021). Using manually curated trait dictionaries, as well as subsets of manually annotated articles, we trained NLP models to perform labelling of entities (taxa, traits, values) and relationships (taxon to trait, trait to value) of thousands of articles. We formally evaluated the performance of our approaches, demonstrating their application to 2’000 publications, which produced 656’403 entity and 339’463 relationship annotations, and highlighting several important technical challenges. Finally, we developed an interactive web tool that makes the results available to the scientific community in the form of the queryable database resource, the ArTraDB Arthropod Trait Database. Together, these tools and resources serve to advance the use of literature mining approaches in biology, ecology, and evolution research, by semi-automating the building of comprehensive open databases of organismal and ecological traits extracted from the literature.

## Materials & Methods

### Sourcing and Processing the Text Corpora

#### Articles Sourced from PubMedCentral via TreatmentBank

To take advantage of the normally highly structured and detailed species information found in taxonomic treatment texts, and at the same time to reduce the complexity of the overall labelling process, the initial corpus of texts was defined using all arthropod species taxonomic treatment texts available from Plazi’s TreatmentBank (Guidoti et al. 2021). Taxonomic treatments refer to sections in scientific publications where the key features describing, distinguishing, and naming a species are documented (Agosti et al. 2022). Treatments have been the building blocks of how data about taxa are provided ever since the beginning of modern taxonomy, and usually follow a highly structured format. This simplifies the first task of labelling taxa (species) because each text already pre-processed by Plazi is directly linked to a known species, meaning at least one annotated taxon in the document should match the linked species name provided by TreatmentBank. From the ∼310’000 treatment texts sourced from Plazi’s TreatmentBank, ∼250’000 were linked to digital object identifiers (DOIs) comprising ∼24’000 unique publications, 3’650 of which were linked with PubMedCentral (PMC) identifiers and thereby presented publicly accessible texts that could be used for labelling and subsequent mining. Note that publications may, and often do, contain many treatments, *i.e.* descriptions of many species, hence the tenfold higher number of treatments compared to the number of publications.

### Processing of PubMedCentral Articles

The PMC article files were retrieved in Extensible Markup Language (XML) format and subsequently transformed into plain text format, maintaining the original text extraction without further modifications such as lowercasing. For the Named Entity Recognition (NER) task, these PMC text files were subsequently converted into the CoNLL format (Kim Sang and De Meulder 2003) (a text file with one word per line with sentences separated by an empty line) using the IOB2 tagging scheme (Inside–Outside–Beginning) (Ramshaw and Marcus 1999). For the Relationship Extraction (RE) task, the same PMC text files were processed into a specialised JSON (JavaScript Object Notation) file format compatible with the “Language Understanding with Knowledge-based Embeddings” model (LUKE) (Yamada et al. 2020). This format splits the text up in context windows, which by default encompass six sentences, along with the offsets and labels for both the head and tail entities.

### Sourcing and Processing Taxonomy and Trait Data

#### Taxonomy Data Sourced from the Catalogue of Life

The Catalogue of Life (COL) represents an authoritative source of taxonomic data built and maintained through a long-term international collaboration of taxonomists and informaticians (Bánki et al. 2024). The COL was therefore selected as the reference taxonomy dataset for building a dictionary of arthropod taxa and for filtering the treatments to select only arthropod species for downstream processing. The monthly COL release of July 2022 (COL Version: 2022-06-23) was processed to extract all accepted taxa (dwc:taxonomicStatus == ‘accepted’) that are hierarchically below Arthropoda (dwc:taxonID == ‘RT’). The dictionary of arthropod taxa therefore contains all accepted arthropod species names along with the taxonomic lineage names ascending the species-genus-family-order-class hierarchy up to the phylum level of Arthropoda. This resulted in a dictionary containing a total of 1’015’642 species and 118’008 higher-level taxonomic names, for use as the input for downstream NER steps to label taxa identified in the processed documents. The COL processing scripts are available as part of the ATResourceManager Snakemake workflow (https://github.com/IDSIA-NLP/ATResourceManager).

### Organismal and Ecological Trait Data

No single, comprehensive, standardised, and machine operable ontology of organismal and ecological traits was available to use to build a dictionary of arthropod trait data. Therefore, extensive manual curation of traits defined across several different resources was required to be as comprehensive as possible while leveraging existing resources and standards. Trait libraries were developed for three broad categories covering arthropod feeding ecology, habitat, and morphology, always requiring that the trait was defined and/or described in an existing online resource. The resources queried included: the Encyclopedia of Life (EOL) (Parr et al. 2014); the Environment Ontology (ENVO) (Buttigieg et al. 2016); the Relation Ontology (RO) (Mungall et al. 2023); the UBERON Anatomy Ontology (Mungall et al. 2012); the BRENDA Tissue Ontology (BTO) (Chang et al. 2021); as well as Wikidata, Wikipedia, and Wiktionary for additional relevant terms with Uniform Resource Identifiers (URIs) that were not incorporated into a formal ontology. Trait names and definitions were inherited from the source ontologies/URIs. Traits were classified into types: “yes/no”, a taxon exhibits or does not exhibit the trait; “association”, one taxon is associated with another taxon through the trait; “measurement” for mass-related traits; “length/width” for measurable body parts; “count” for countable body parts. Synonyms of trait names were automatically generated by scraping synonyms from Synonyms.com and enhanced with word vectors from PubMed and the Common Core to obtain related terms. This approach was refined by implementing an improved search for synonyms, creating a new table that included only those synonyms appearing at least ten times in the 5’000 articles from the ZooKeys journal that were available in PMC. This method was devised to capture real words commonly used in taxonomic treatments, managing to generate synonyms for most terms. However, the synonyms contained instances of inaccurate or inappropriate terms so manual curation was applied to reduce redundancy and improve informativeness of the alternative phrasings including pluralisations. This resulted in a dictionary containing a total of 390 traits (81 feeding ecology; 184 habitat; 125 morphology, Supplementary File S1), to be used as the input for downstream NER steps to label traits identified in the processed documents.

### Curating Gold-Standard Annotation Data

To fine-tune and formally evaluate the performance of the NLP models employed for the NER and RE tasks (see below), two entomology domain experts annotated a set of 25 articles randomly selected from the 3’650 obtained from PMC. The two annotators employed the tagtog text annotation tool (Cejuela et al. 2014) that provides a user-friendly interface to manually annotate and normalise entities in documents imported from PMC, as well as to add entity labels, relationships, and more. The annotators worked independently on their assigned documents to avoid biasing each other, however, they did develop a set of guidelines during the annotation process to describe the steps to follow for dealing with some more complex cases (Supplementary File S2). For example, in addition to the required entities “taxon”, “trait”, and “value”, and the required relationships “has_trait” and “has_value”, the entity “qualifier” and relationship “has_qualifier” were created in order to label and link taxonomic qualifiers to labelled taxa entities such as “female”, “male”, “juvenile”, or “larva”. To be able to compare annotator styles, a subset of five documents was independently annotated by both experts. The completed annotations of the 25 documents were exported from tagtog. Each document, originally in tagtog’s native HTML format accompanied by annotations in JSON format, was converted into a BioC JSON file (Comeau et al. 2013). This format serves as the base for all subsequent processes. The documents annotated by both experts were used to calculate the inter-annotator agreement by means of Cohen’s Kappa score (Cohen 1960). The agreement was evaluated by comparing exact and partial matches of named entities, as well as exact matches of relationships within a tolerance of 4 characters for the entity offset boundary in these documents. To be able to formally evaluate the performance of the NLP models the gold-standard annotation data were randomly split into a training (TRAIN-GOLD) and an evaluation (TEST-GOLD) dataset (Supplementary File S3). The TRAIN-GOLD dataset contained four documents (812 entities, 641 relationships) and the TEST-GOLD dataset contained 21 documents (4’272 entities, 3’439 relationships), corresponding to a 1’453/7’711 (18.8%) ratio between TRAIN-GOLD and TEST-GOLD. The data split was deliberately chosen, opting for a larger TEST-GOLD dataset to ensure it is statistically robust for valid analysis. Simultaneously, the TRAIN-GOLD dataset was intentionally kept small enough to train NER and RE models under a low-resource setup.

### Natural Language Processing for Entity and Relationship Annotation

For automatic identification of arthropod trait value triples, a two-step pipeline was developed: First, entities (arthropods, traits, and values) are recognised (identified in the texts) and normalised (mapped to dictionaries of known entities, if possible) and then the underlying relationships between arthropods and traits and between traits and values are extracted (annotated) where possible. Different entity recognition systems were therefore implemented to automatically recognise arthropods (taxa), traits, and values, and then normalise the recognised entities according to a set of terminologies. To identify the relationships between the recognised entities, a relationship extraction model was applied where each relationship connects exactly two entities and the relationship type is determined by the types of connected entities.

#### Named Entity Recognition (NER)

The NER steps employed and tested several Bidirectional Encoder Representations from Transformers (BERT)-based models, namely BERT-large-uncased (Devlin et al. 2019), RoBERTa-large (Liu et al. 2019), and the domain-adapted BioBERT (Lee et al. 2020) model initialised on the BERT model and further pre-trained on PubMed abstracts and PMC complete articles. For NER of value entities, pre-trained models were used from SpaCy (Montani et al. 2023) and quantulum3 (Mündler 2024), which are libraries that are specialised in flexible matching of measurements and their entities.

#### Named Entity Normalisation (Linking) (NEN)

To perform NEN, the OntoGene BioMedical Entity Recogniser (OGER) was used, OGER offers a set of tools for text mining and information extraction (Basaldella et al. 2017, Furrer et al. 2022). The normalisation (or linkage) of entities is achieved by flexible matching of the recognised entities with the curated arthropod and trait dictionaries described above.

#### Relationship Extraction (RE)

The Transformer-based LUKE (Yamada et al. 2020) model was used for the RE task. The LUKE model takes a text string along with the offsets of a head and tail entity to perform classification according to a set of relationship labels.

### Technical Specifications of the ArTraDB Web Resource

ArTraDB is a web application designed and built to present the predicted annotations to the scientific community. The following technology stack was used to build the resource: data are stored in a Neo4J database (https://neo4j.com/) and made available through a backend Application Programming Interface (API) based on express.js (https://expressjs.com); the frontend was built using Vue.js (https://vuejs.org) and node.js (https://nodejs.org); for visualisation of annotated documents the TextAE annotation editor (https://textae.pubannotation.org) (Lever et al. 2020) was integrated into the web application.

## Results

### A Workflow for Annotating Arthropod Organismal and Ecological Traits

The analytical workflow for processing and annotating thousands of articles to identify organismal and ecological traits of arthropods (Figure 1) consists of several key data preparation steps (ATResourceManager and Domain Expert curation) and model training procedures (ATTrainer), in order to subsequently perform the text mining tasks (ATMiner) to produce the predictions and upload them for viewing in ArTraDB. Firstly, the domain expert curation tasks resulted in two key outputs: the Gold Standard Annotations and the Curated Trait Vocabularies. A set of selected documents was manually annotated by domain experts to provide a resource for downstream training and for assessing the performance of the text mining tasks (see Methods). The domain experts also built curated trait vocabularies (including synonyms) covering the three categories of feeding ecology (n=81), habitat (n=184), and morphology (n=125), based on combinations of existing ontologies and online resources (see Methods). In parallel, the ATResourceManager preparation steps were developed to: (1) process the taxonomic treatment documents from Plazi and retrieve the corresponding publications from PMC; (2) extract from the Catalogue of Life taxonomy all accepted arthropod species and their higher-level taxonomic names; and (3) extract from the Encyclopedia of Life traits database all available taxon-trait annotations for arthropods (see Methods for details). Subsequently, the ATTrainer language model training steps take as input the Gold Standard Annotations (TRAIN-GOLD subset) for the fine-tuning of the BioBERT model and for the training of the LUKE model (see Methods). These models are then used in the ATMiner tasks for Named Entity Recognition (NER) with BioBERT and Relationship Extraction (RE) with LUKE, also using the curated trait vocabularies to perform entity normalisations using OGER (see Methods). The resulting predicted annotations - the entities of arthropods, traits, and values - and the arthropod-trait and trait-value relationships were then imported into the ArTraDB web resource where they can be reviewed by the community.

**Figure 1:**
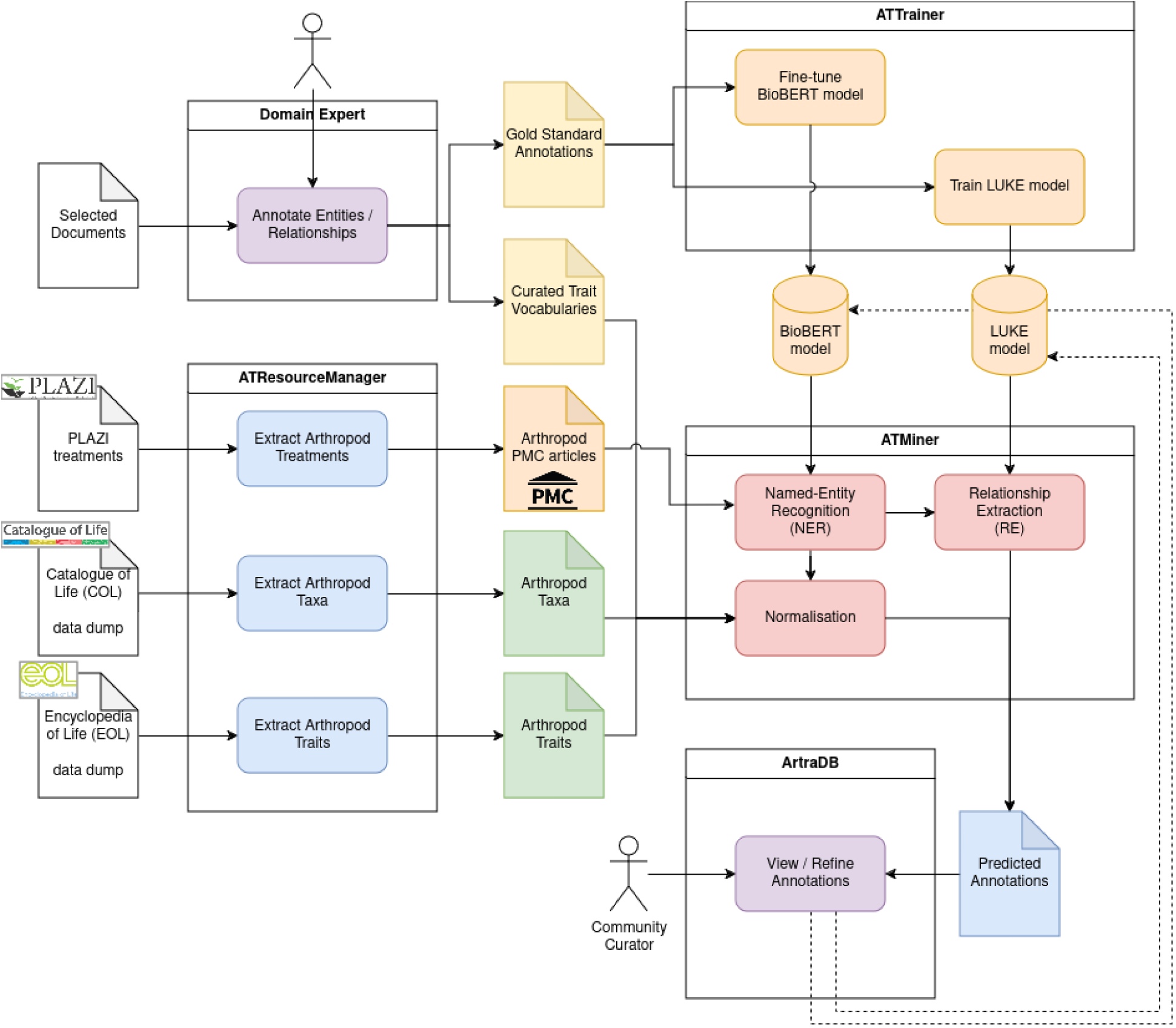
The arthropod organismal and ecological traits annotation workflow. The workflow starts with curation performed by domain experts resulting in entity and relationship annotations for a selected subset of publications as well as curated vocabularies of sets of organismal and ecological traits. The ATResourceManager steps include the processing of data sourced from the Catalogue of Life (taxonomy) and the Encyclopedia of Life (arthropod-trait relationships) to generate taxon and trait dictionaries, as well as the retrieval of publications for processing from PubMed Central based on the selection of all Plazi TreatmentBank records for arthropods. The expert-generated Gold Standard Annotations are used as input to train (ATTrainer steps) Natural Language Processing (NLP) models for the Named Entity Recognition (NER) and Relationship Extraction (RE) tasks, with the trait vocabularies and taxa dictionaries being used for entity normalisation (ATMiner steps). Finally, the predicted annotations are made available to the scientific community via the ArTraDB web resourc where community curators could potentially provide corrections to the annotations that can later be used for refinement of the NER and ER models (dotted lines).

### Entity and Relationship Annotation of PubMedCentral Articles

#### Annotation Results for Entity and Relationship Discovery

The application of the workflow presented in Figure 1 to a total of 2’000 publications sourced from PMC resulted in the annotation of 656’403 entities (arthropods, traits, and values) and 339’463 relationships (hasTrait, hasValue), summarised in Figure 2. The PMC articles range in lengths from 173 to 27’466 characters with a median of 15’720 and an interquartile range from 11’452 to 20’506 (Figure 2A). The densities of entity and relationship predictions are highest for entities of type “value” and hasValue relationships, with medians of ∼10 annotations per 1’000 characters, arthropod and trait entities have median densities of four and six annotations per 1’000 characters respectively, and hasTrait relationships has the lowest density with a median of zero annotations per 1’000 characters (Figure 2B). In contrast, the 25 articles comprising the gold standard annotation dataset show median densities of 4.9, 6.4, and 6.1 annotations per 1’000 characters for arthropod, trait, and value entities, respectively, and the hasTrait and hasValue relationships show medians of 6.4 and 6.1 annotations, respectively (Figure 2C). These manually annotated documents (17 of which were complete articles and eight only abstracts) contained in total 4’990 named entities (1’069 arthropods, 2’078 traits, and 1’843 values) and 3’628 relationships (1’777 hasTrait and 1’851 hasValue). For the predicted annotations, the total numbers of entities and relationships are generally higher in longer documents, reaching over 900 and over 600 annotations, respectively (Figure 2D). This trend is replicated when considering each entity and relationship subtype separately, with the largest numbers of annotations identified in some of the longest documents, reaching maxima of 456, 393, and 669 for arthropods, traits, and values, and 100 and 717 for hasTrait and hasValue, respectively (Figure 2E). At minimum, a publication should contain one taxonomic treatment describing a single arthropod species, e.g. “*Pachybrachis sassii*, a new species from the Mediterranean Giglio Island (Italy) (Coleoptera, Chrysomelidae, Cryptocephalinae)” (Montagna 2011) (length: 13’550 characters; annotated entities: 63 arthropod, 79 trait, 149 value). However, the much longer articles generally describe a whole group of species for a particular region, *e.g.* “The dipteran family Celyphidae in the New World, with discussion of and key to world genera (Insecta, Diptera)” (length: 27’381 characters; annotated entities: 172 arthropod, 141 trait, 246 value) contains 92 taxonomic treatments (Gaimari 2017).

**Figure 2:**
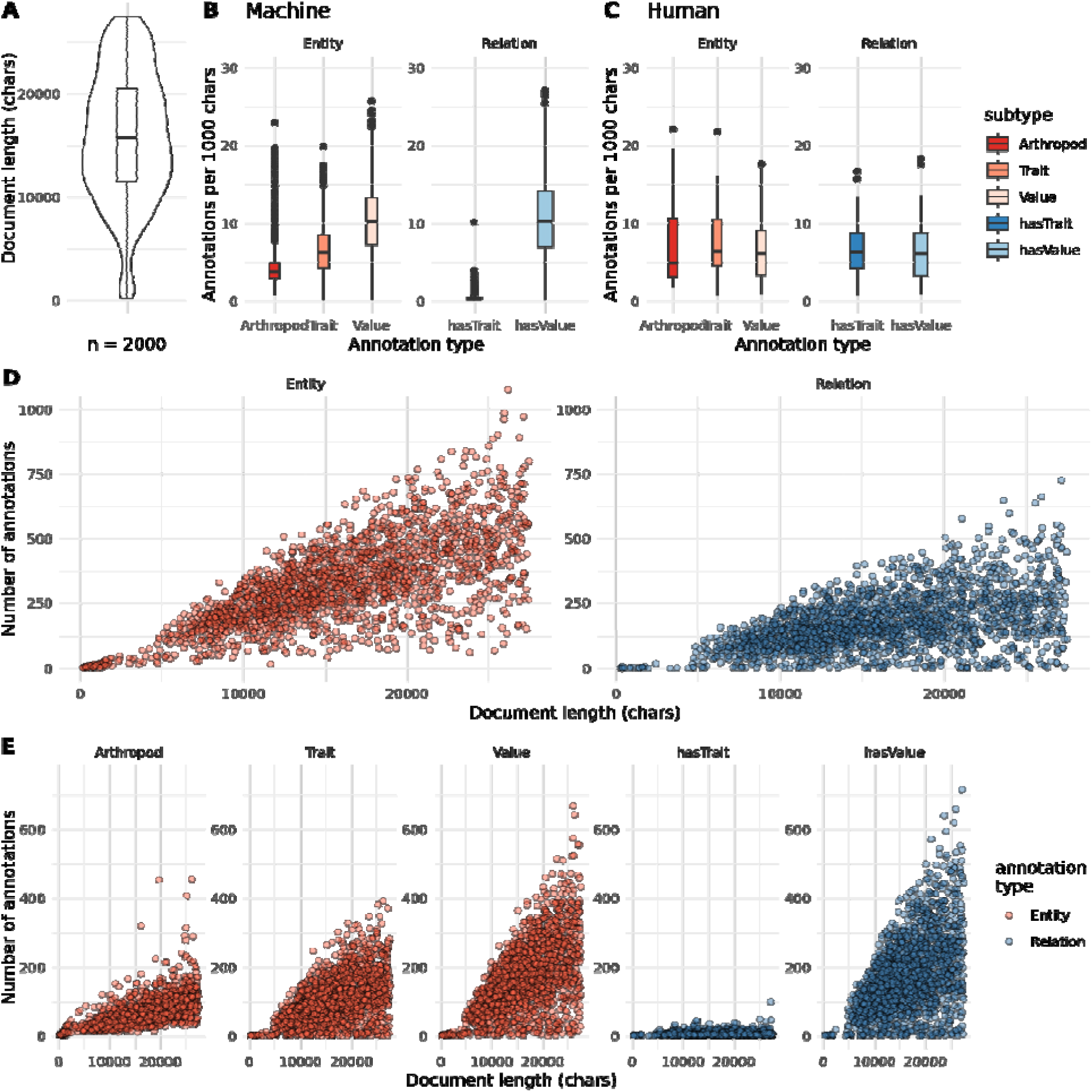
Distributions of PMC article properties and the resulting entity and relationship annotations. (**A**) The input dataset consisted of 2’000 PMC articles exhibiting a broad character (chars) length distribution. The relative number of resulting predictions of annotated entities (arthropods, traits, values) and relationships (taxon to trait - hasTrait, trait t value - hasValue) are shown for the whole dataset (**B**) and for the 25 gold-standard manually annotated documents (**C**). The absolute number of predicted entity and relationship annotations compared to the document lengths in characters is shown for annotation types (**D**) and subtypes (**E**). Boxplots in panels **A, B**, and **C** show the median, first and third quartiles, and lower and upper extremes of the distribution (1.5 × Interquartile range).

### Assessing the Complexity of the Task by Examining Inter-Annotator Agreement

To begin to interpret the prediction results from the workflow it is important to understand the complexity of the annotation task itself, insights into which can be gained by examining the levels of agreement between the curated annotations generated by the two domain experts. The five documents that were annotated by both domain experts included two complete publications and three abstract-only articles (Figure 3). In total for these five documents, the two experts annotated 1’477 named entities (161 arthropods, 764 traits, 552 values), with annotator 1 identifying 80 arthropods, 416 traits, and 334 values, and with annotator 2 identifying 81 arthropods, 348 traits, and 218 values. They also annotated a total of 1’094 relationships (553 hasTait, 541 hasValue), with annotator 1 identifying 343 hasTrait and 343 hasValue relationships, and with annotator 2 identifying 210 hasTrait and 198 hasValue relationships. Cohen’s kappa is used to measure inter-annotator reliability, or concordance, to assess the degree of agreement amongst independent observers who rate, label, or classify the same phenomenon, with values below 0.6 generally indicating inadequate agreement (McHugh 2012). Amongst the five documents curated by both annotators, Cohen’s kappa values reflect varying levels of inter-annotator agreement for entities (Figure 3B), from poor agreement (∼0.35), to moderate agreement (∼0.5), to substantial agreement (∼0.8), with lower agreement levels for relationships (Figure 3C). While Cohen’s kappa scores provide a standardised measure of agreement, their values must be carefully interpreted within the study’s context, where here they serve to highlight the complexity of the annotation tasks.

**Figure 3:**
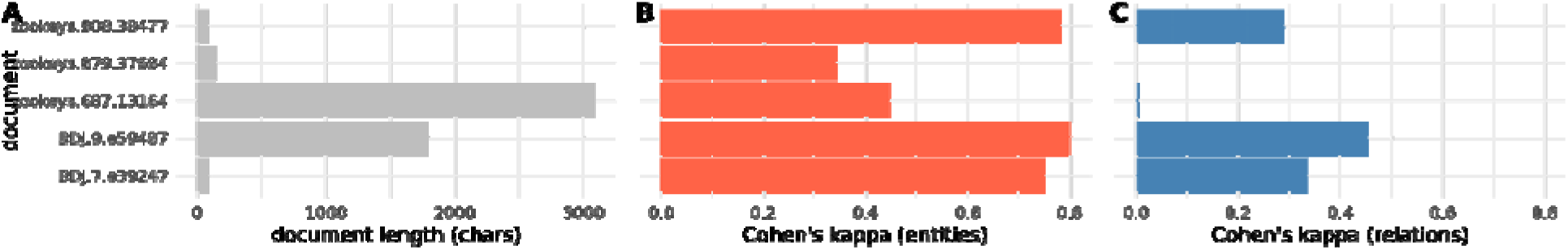
Inter-annotator agreement of five documents annotated by both experts. (**A**) The bars show document lengths in characters (chars). (**B**) The bars show the level of annotation agreement for entities (taxon, trait, or value) between the two annotators as measured using Cohen’s kappa. (**C**) The bars show the level of annotation agreement for relations (hasTrait or hasValue) between the two annotators as measured using Cohen’s kappa.

### Assessing Entity Normalisation with the Taxon and Trait Dictionaries

Entity normalisation, or linking, which was performed using OGER, is the process of matching the entities that were annotated in the articles with the dictionaries of arthropods (taxa from the Catalogue of Life) and traits (the collated sets of feeding ecology, habitat, and morphology traits), with the goal of assigning to each labelled entity a unique identifier from one of the input resources. It is important to understand the performance of the normalisation task in order to interpret the quality of the entity prediction results from the workflow. The taxon dictionary contained a total of 1’015’642 arthropod species and 118’008 higher-level taxonomic names and the trait dictionary contained a total of 390 traits: 81 feeding ecology; 184 habitat; 125 morphology (see Methods). Focusing on the identification and quantification of taxon and trait entities within the article corpus, the coverage and frequency of entities mapped to the predefined dictionaries and those that could not be mapped provide an assessment of the entity normalisation process and the comprehensiveness and relevance of the dictionaries (Figure 4).

**Figure 4:**
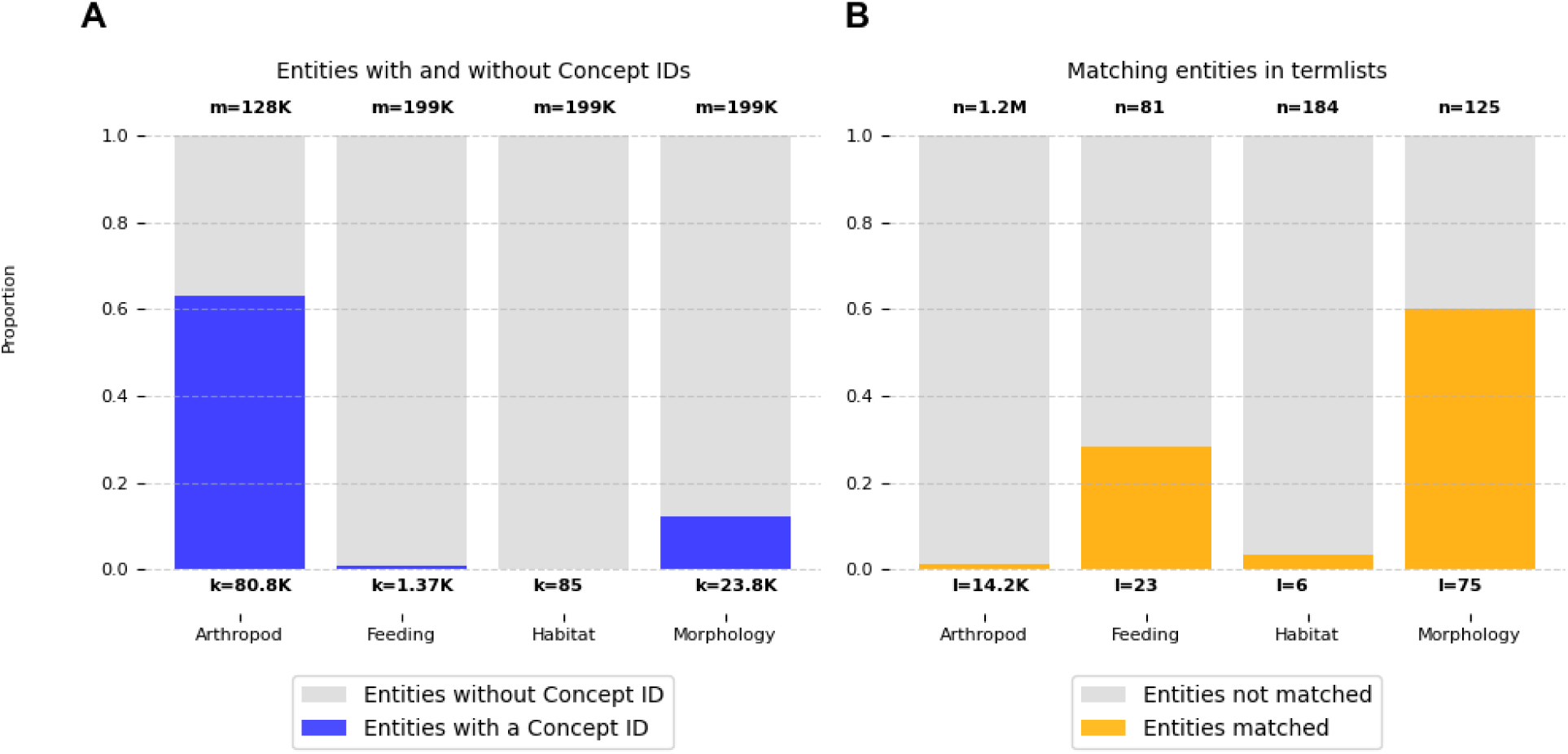
Taxon and trait dictionaries compared with annotated entities. For the 2’000 PMC articles analysed: the proportions of all annotated entities that could be mapped to the corresponding “Concept IDs” of the taxon and trait dictionaries (**A**), and the proportions of taxon and trait dictionary terms that were matched with annotated entities in any article (**B**). In (A) ‘m’ represents the total numbers of taxon and trait entities and ‘k’ indicates how many of these were mapped to Concept IDs in the dictionary termlists (for trait entities divided into feeding ecology, habitat, and morphology).In (B) ‘n’ represents the total numbers of taxa and traits in the dictionary termlists and ‘l’ indicates how many of these were matched in the articles.

Across the 2’000 articles processed by the workflow, a total of 128’149 taxon entities were annotated with 63% matching terms in the dictionary (mapped to Concept IDs), comprising 24’207 species and 56’532 higher-level taxonomic names (Figure 3A). Notably, taxa such as the order Hymenoptera (sawflies, wasps, bees, and ants), or the genus Tipula (crane flies), were amongst the most frequently annotated entities, with 388 and 384 occurrences, respectively. Reviewing some of the 47’312 non-mapped taxon entities revealed examples of correctly identified arthropod taxa which nevertheless are not included in the Catalogue of Life and are therefore not in the dictionary, e.g. the genera Micrencaustes (beetles) and Deinodryinus (parasitoid wasps). Of the 199’276 (28’348 unique) trait entities annotated, 12.7% were successfully mapped to the trait dictionary (linked to Concept IDs), with 1’366, 85, and 23’816 entities mapping to feeding ecology, habitat, and morphology terms, respectively (Figure 3A). Note that the NER task labels entities as taxa, traits, or values, further categorisation of traits into feeding ecology, habitat, and morphology terms is only possible when entity normalisation is successful. Feeding and morphology traits such as “host” (273), “mouth” (16), and “legs” (1’663) were amongst the most prevalent annotated entities. Many annotated trait entities could not be mapped (168’237), one of the most frequent being “distribution”, with 2’506 occurrences. Considering instead the numbers of unique terms in the dictionaries that could be annotated and linked in the articles, only 14’243 of the 1.2 million arthropods in the dictionary (1.2%) were identified, and from the trait dictionary of feeding ecology, habitat, and morphology terms, 28.4%, 3.3%, and 60%, respectively, were annotated and linked (Figure 3B). Failure to identify dictionary terms in the annotated articles may be because the terms are simply not present (it cannot be expected that a small subset of 2’000 articles will contain mentions of all 1.2 million described arthropod species), or because normalisation was unable to link terms and phrases recognised as entities in the texts with the terms, phrases, and synonyms that make up the concepts of the dictionaries.

### Performance Comparisons of Natural Language Processing Models

#### Named Entity Recognition Baseline Performance

An evaluation of a Named Entity Recognition (NER) baseline was conducted across various configurations. Several general and domain-specific pre-trained language models were fine-tuned on the TRAIN-GOLD dataset. To train the models, the dataset was converted to IOB2 format. Two evaluation methods were employed for the results presented in Figure 5: the Conference on Natural Language Learning (CoNLL) evaluation and strict metrics. The reported results are based on the F1-score (F) and corresponding Precision (P) and Recall (R). Under the CoNLL evaluation, the baseline demonstrated a macro-average 0.56 F (0.55 P / 0.57 R) and a weighted-average of 0.52 F (0.53 P / 0.53 R) across all entity types. Notably, entities classified as ‘Arthropod’ achieved the highest F1-score at 0.74 F (0.7 P / 0.78 R), signifying superior recognition capabilities in comparison to other categories. Conversely, ‘Value’ entities posed greater challenges, with the lowest score of 0.37 F(0.32 P / 0.43 R). This indicates substantial difficulties in the precise identification of these entities. ‘Value’ entities encompassed a diverse array of concepts, ranging from measurements (*e.g.*, ‘56.6 mm’) and colour descriptors (*e.g.*, ‘brownish-yellow’) to locations (*e.g.*, ‘China’). This disparity highlights the model’s varied performance across different entity types. When evaluated using the strict metric, a notable enhancement in both precision and F1-scores was observed for most entity types, compared to the CoNLL evaluation metric (Figure 5). ‘Arthropod’ entities maintained the highest score 0.78 F (0.78 P / 0.78 R), consistent with the previous evaluation. The overall macro- and weighted-average scores increased to 0.59 F (0.63 P / 0.57 R) and 0.56 F (0.6 P / 0.52 R), respectively, indicating a more accurate entity recognition when the strict metric was applied. This comparison not only underscores the baseline’s strengths and weaknesses in recognising various entities but also highlights the impact of evaluation criteria on perceived performance.

**Figure 5:**
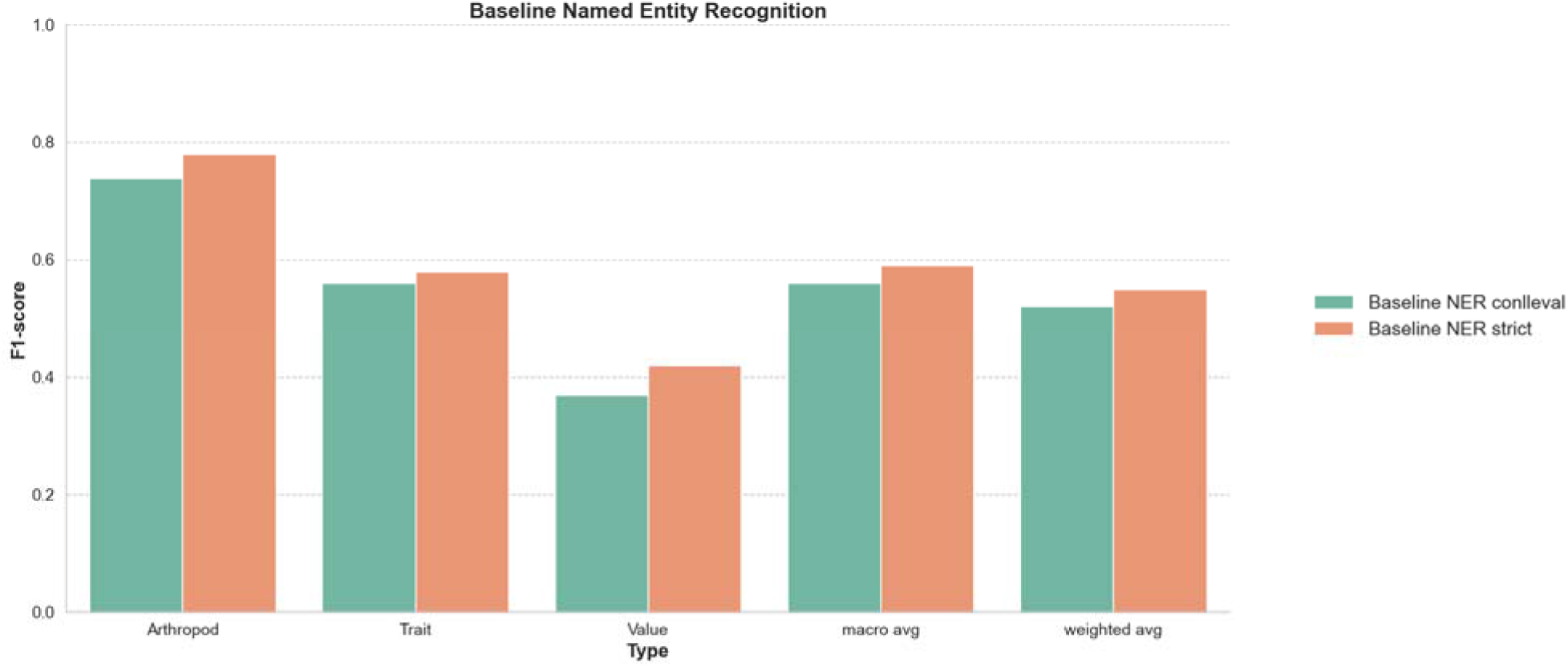
CoNLL evaluation and strict F1-score baseline results for the named entity recognition. The F1-score performance of the baseline Named Entity Recognition (NER) model on the test set of PubMedCentral (PMC) articles using the Conference on Natural Language Learning (CoNLL) evaluation (conlleval) and strict methods. The scores are shown explicitly for three entity types: Arthropod, Trait, and Value. Additionally, for the three types combined, the macro-average F1-score, which does not account for class imbalances, and the weighted average F1-score, which adjusts for the imbalance of different classes, are also presented. A detailed listing of all baseline results can be found in Supplementary File S4.

### Relationship Extraction Baseline Performance

Figure 6 outlines the outcomes of the Relationship Extraction (RE) baseline across three different configurations of the LUKE model, namely “NCB” (None-Class Balanced), which limits the amount of ‘none’ relationships during training to match the majority class, “Tag” which uses XML to tag the entities inline, and “Long-Range”, which captures long-range relationships via a different training setup. By default the LUKE model was used with a shifting context window spanning 1 to 6 consecutive sentences to detect relationships. In contrast, for the Long-range approach a version of the LUKE model was fine-tuned by extracting and merging the two target entities with their 500 surrounding characters each. The NCB approach, even after balancing the frequency of ‘none’ classes with the most common relationship (‘hasValue’), continued to face challenges in accurately identifying specific relationships like ‘hasTrait’. This indicates persistent difficulties in detecting nuanced or less common relationships. Furthermore, applying the Tag approach in addition to the NCB approach improved the RE baseline in the macro-average score from 0.57 F (0.62 P / 0.66 R) to 0.65 F (0.66 P / 0.69 R) compared to the standard NCB configuration. This suggests that entity tag information contributes positively to relationship extraction performance. Using NCB and Tag combined with the Long-range RE setup demonstrated an interesting pattern with ‘hasValue’ relationships, where a perfect recall (1.00) but very low precision (0.02) resulted in a low F1-score (0.03). This indicates the model’s tendency to over-identify ‘hasValue’ instances, leading to numerous false positives. The prediction results for the ‘hasTrait’ relationship show 0.2 F (0.13 P / 0.52 R) while scoring for ‘none’ relationship 0.92 F (1.0 P / 0.86 R), and a macro-average of 0.38 F (0.38 P / 0.79 R) , making it the worst-performing baseline.

**Figure 6:**
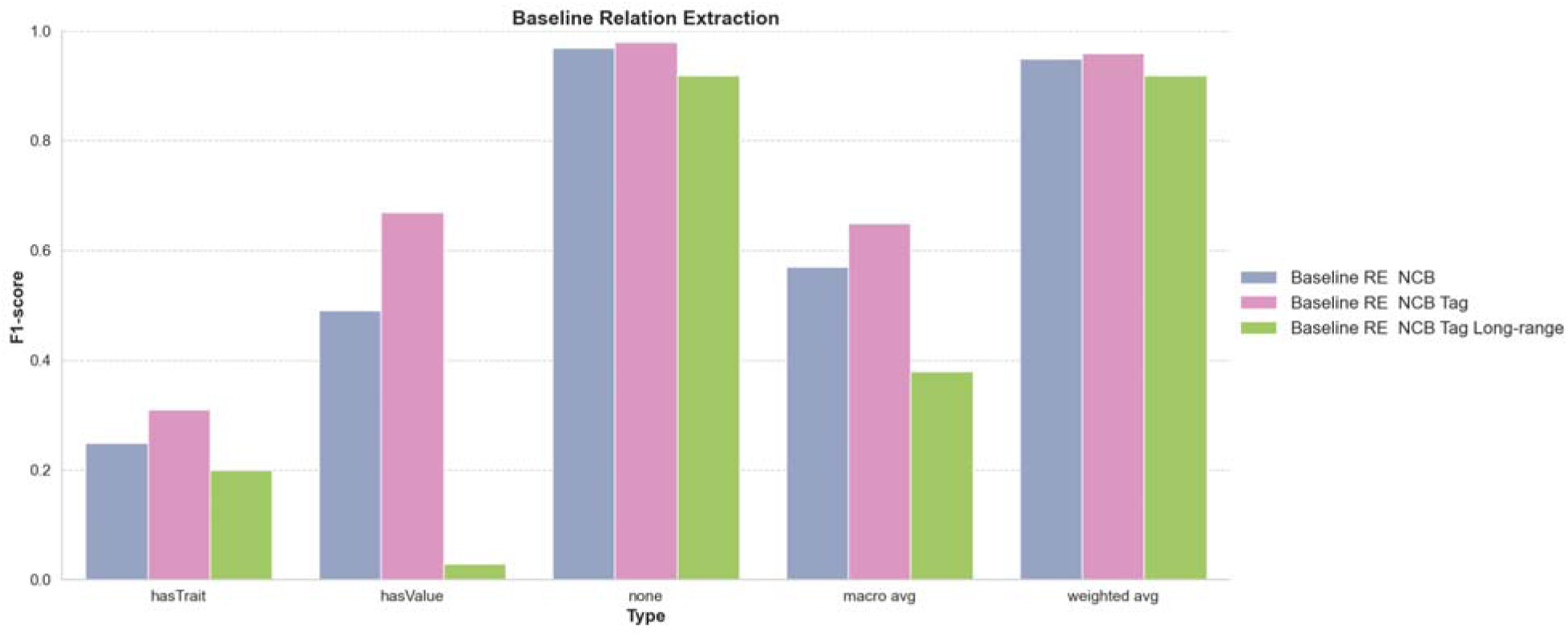
NONE-class balanced (NCB), entity tagged (Tag), and long range baseline results for relationship extraction. The F1-score performance of three baseline configurations of the Relationship Extraction (RE) model on a test set of PubMedCentral (PMC) articles. The configurations are: Non-Class-Balanced (NCB), which limits the amount of ‘none’ relationships during training to match the majority class; XML Inline Tag Entities (Tag); and Long-range Context, which provides a context of 250 characters around each target entity rather than using a sliding context window. Performance scores are specifically shown for two relationship types: ‘hasTrait’ between Arthropod and Trait, ‘hasValue’ between Trait and Value, and ‘none,’ which indicates the absence of a relationship. Additionally, the figure includes both the macro average F1-score, which does not account for class imbalances, and the weighted average F1-score, which compensates for these imbalances. A detailed listing of all baseline results can be found in Supplementary Table S4.

### The Arthropod Trait Database ArTraDB Web Resource

The annotation predictions obtained from applying the workflow to the PMC articles are made available to the community through the dedicated web application, ArTraDB: the Arthropod Trait Database (https://artradb.unil.ch). The results are presented in a simple table-like view where each row represents a single entity annotation, pairs of entities connected by either a hasTrait or hasValue relationship, or complete trio annotations of connected Arthropod-Trait-Value entities (Figure 7). The ArTraDB resource was designed and developed to provide two main functionalities: (i) Browse and search facilities enabling the identification of predicted species/taxa and/or traits and/or values within the set of annotated documents; and (ii) Browsable visual displays of the predicted entity and relationship annotations in the local context of the corresponding document.

**Figure 7:**
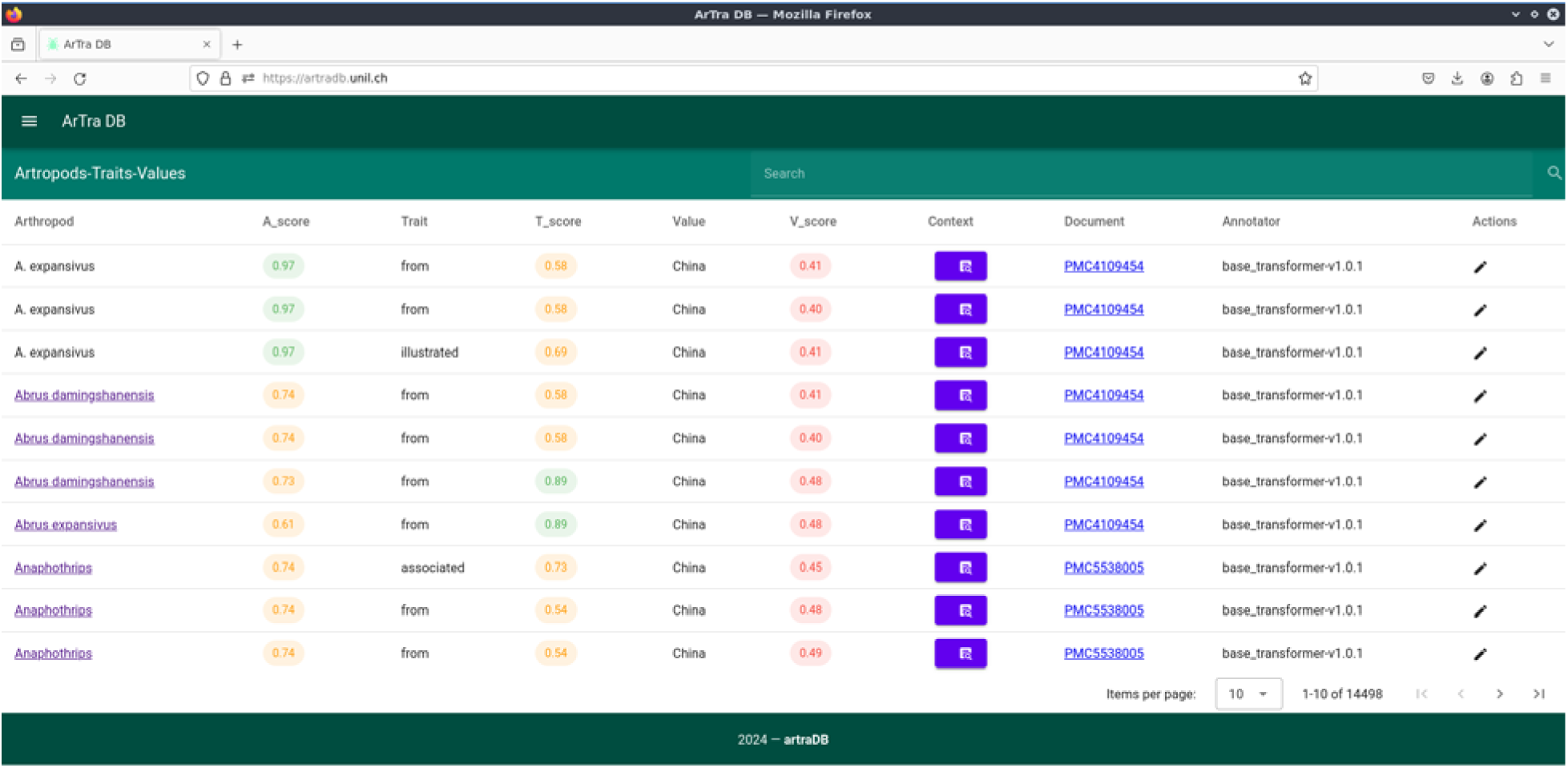
ArTraDB table view browse and search functionalities. The table view of annotations displays rows of annotated entities with their corresponding Named Entity Recognition (NER) confidence scores, a clickable icon to open the Annotation Viewer window, the hyperlinked PubMedCentral (PMC) article identifier, and the source of the annotation, in a browsable paginated format with a user-configurable number of items to display per page. Annotated arthropods and traits that were successfully linked to Concept IDs in the corresponding dictionaries are hyperlinked to the corresponding source definition, *e.g.* the Catalogue of Life (COL) for arthropods, and the Encyclopedia of Life (EOL) or other resources for trait entities. Indexing of the ArTraDB data allows for rapid user searches to filter the complete table to rows with entries matching terms entered in the simple search box above the table.

### ArTraDB Browse and Search Functionalities

The browsable table view of the workflow-predicted annotations for the set of processed articles provides a paginated display of rows of annotated entities together with the NER confidence scores assigned by the prediction algorithm (Figure 7). Where both hasTrait and hasValue relationships connect an arthropod entity with a trait entity and that same trait entity to a value entity the row represents a complete Arthropod-Trait-Value trio annotation. When either hasTrait or hasValue relationships are lacking, the row displays only the Arthropod-Trait or Trait-Value pairs. If no relationships were predicted, the arthropod and trait entities are displayed as single annotations with their corresponding scores, while the value entities are omitted. When entity normalisation was able to successfully link annotated arthropods and traits to Concept IDs in the corresponding dictionaries, these are hyperlinked to the corresponding source definition, e.g. the Catalogue of Life for arthropod entities, and the Encyclopedia of Life or ontology and Wiki resources for trait entities. Additional columns in the table view include a clickable icon to open the popup Annotation Viewer window (Figure 8), the PMC identifier hyperlinked to the fully annotated document, the source of the annotation (*i.e.* version of the workflow that made the predictions). The table view can be browsed page by page with a user-configurable number of rows to display per page. The data in each column are indexed to enable rapid user searches using the simple search box above the table that filters the results to contain only rows matching the entered search term. This simple table view of the annotations provides an intuitive browsable and searchable interface to the thousands of annotations produced by the workflow.

**Figure 8:**
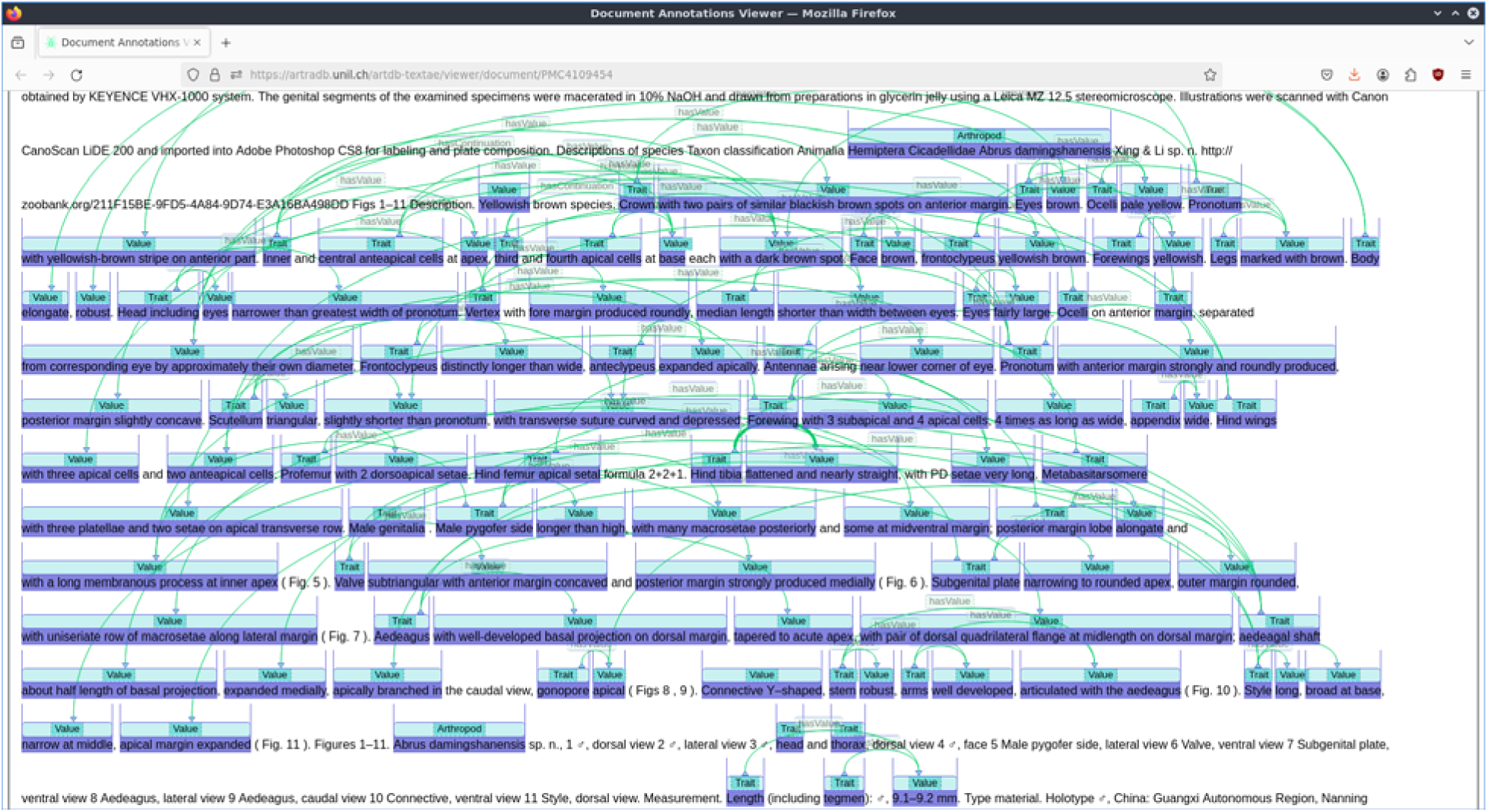
ArTraDB document view for visualising all predicted annotations in an article. Clicking the hyperlinked PubMedCentral (PMC) identifier in the ArTraDB table view opens a document view of the fully annotated article showing all entities and relationships. The entities are highlighted and labelled as Arthropod, Trait, or Value entities, with the predicted relationships indicated by lines connecting the relevant entities. This provides users with the full-text context of each predicted entity or relationship annotation within the document.

### ArTraDB Document Context View of Annotations

For each row of predicted entity or entity and relationship annotations, a clickable icon in the Context column allows users to open an annotation viewer window that shows these labelled entities and any predicted relationships in the local context of the source text. Alternatively, clicking the hyperlinked PMC identifier opens a document view of the fully annotated article showing all entities and relationships. In both viewers, the entities are highlighted and labelled as Arthropod, Trait, or Value entities, with the predicted relationships indicated by lines connecting the relevant entities (Figure 8). These visualisation functionalities allow users to view the predictions in their local and global contexts, to be able to manually assess the reliability of the automatically generated annotations.

## Discussion

Focusing on the methodological explorations of the opportunities presented by modern literature mining approaches, we aimed to annotate available arthropod organismal and ecological trait data to facilitate the automated extraction of knowledge from publications. This involved building and testing an analytical workflow for processing and annotating thousands of PMC articles containing taxonomic treatments of arthropods. It also required the collation of curated trait vocabularies and the development of model training procedures to perform the text mining tasks and formally test the performance of various approaches. Finally, the resulting predicted annotations of the entities of arthropods, traits, and values, as well as the arthropod-trait and trait-value relationships were made available for community review through the deployment of the open online ArTraDB web resource. Here we discuss the key findings and important challenges encountered while exploring the utility and performance of the different methodologies and approaches we tested to achieve our aims.

### Results are variable because annotation is an inherently complex task

The process used to annotate the articles and train the models presents several methodological challenges that influence the efficiency and accuracy of predicting arthropod-trait-value triples. A key challenge was the generation of training data through the manual annotation of documents by domain experts. Compared to the predictions, the annotators identified higher median densities of arthropod and trait entities and hasTrait relationships, while the automated processes produced higher median densities for value and hasValue annotations (Figure 2). Nevertheless, assessing inter-annotator agreements highlighted the complexity of the annotation tasks and the challenge of uniformity in defining entity boundaries or relationships between entities (Figure 3). Defining entity boundaries and relationships in complex cases are subjective decisions that can diverge based on the annotator’s understanding and experience. For example, “*mesoscutum silvery setae evenly distributed*”: could be simply annotated as trait=“*mesoscutum*” and value=“*silvery setae evenly distributed*”, or by introducing a discontinuous trait “*mesoscutum setae*” with two associated values “*silvery*” and “*evenly distributed*”. The variability in manual annotations introduces conflicts in the input data that serve as the basis for training the NER and RE models, which therefore negatively impacts the model performance. While independence is required to formally assess inter-annotator agreement for such tasks, the recommendation from this work is to build a set of guidelines using examples encountered in the documents. The guidelines represent the consensus of the domain experts on how to proceed technically with annotating complex cases, and therefore serve to enhance consistency and ultimately provide better input data for training the models. Indeed guidelines and annotator training in the context of constructing NER and RE benchmarking corpora of abstracts and metadata files from biodiversity datasets achieved inter-annotator entity agreements of 0.76 and 0.70 for two pairs of annotators (Abdelmageed et al. 2022). As well as the manual annotation variability, the low amount of training data - the 25 articles comprising the gold standard annotation dataset - also posed challenges. One strategy to increase the amount of training data could be to use distant supervision techniques, such as employing large language models (LLMs) to generate structured annotated texts based on the manually-annotated examples (Li et al. 2024). This could introduce biases and would likely be limited to shorter text bodies and smaller annotation context ranges than those encountered in the published articles, but the approach remains worth considering to augment training data. Ultimately, more consistent and better quality annotations across many more articles could be achieved through community review of the NER and RE predictions. The integration of such community-reviewed annotations into future training datasets (Figure 1) could therefore serve to iteratively improve model performance with a growing corpus of articles in the gold standard annotation dataset.

### Entity recognition performs better than relationship extraction

The NER and RE tasks present distinct challenges as evidenced by the performance disparities in the baselines (Figure 5, Figure 6). NER focuses on identifying and classifying entities in the text, with results showing a varied performance across different entity types, where “Arthropod” entities were consistently easier to correctly recognise than “Value” entities and “Trait” entities showing intermediate performance (Figure 5). In contrast, RE aims to identify relationships between entities, which is an inherently more complex task due to the need to understand context and entity interactions where the entities themselves may not be correctly or completely identified. The results indicate that while the baseline is proficient in identifying the absence of a relationship (“none”), it struggles with more specific and low-frequency relationships. The balancing of classes in the training and the introduction of entity tags slightly improved the performance but also revealed the model’s limitations in generalising across different types of relationships (Figure 6). Of particular interest with respect to the processing of taxonomic treatments is the question of context sizes when attempting to identify entity relationships. The hasTrait relationships can often be long-range and few-to-many because the arthropod may be mentioned a few times near the start of an article where the text that follows is then dense with many specific trait entities, *e.g.* a paragraph with detailed descriptions of the morphology of the arthropod. In contrast, the hasValue relationships are more often likely to be short-range and one-to-one as the values of the mentioned traits are usually presented in very close proximity (same sentence). The close proximity can still present substantial challenges, *e.g.* in taxonomic treatments where lists of traits are followed by corresponding lists of values: “*Antennal segments III–VIII length 38, 47, 43, 41, 33, 21*” (here with the added complexity of an inferred list of segments III, IV, V, VI, VII and VIII). Given these differences, alternative strategies where training and RE steps are carried out separately for hasTrait and hasValue relationships might perform better. Overall, the distinction between the NER and RE tasks is evident here in their respective challenges and baseline performances. NER requires accurate classification of individual entities, while RE demands a deeper understanding of the context and the interactions between entities. This difference in complexity was also reflected in the lower inter-annotator agreement scores achieved for RE than for NER annotations (Figure 3), which would have reduced the effectiveness of training the LUKE model. While the use of NER and RE in mining biodiversity literature has shown promising results, the complexities associated with training data quality and annotation consistency remain substantial challenges. Addressing these issues will be essential for advancing the capabilities of NLP applications in biodiversity research.

### Entity normalisation is a critical yet challenging process

The task of normalisation involves linking the identified entities to defined dictionaries, vocabularies, or ontologies with accompanying descriptions and associated information. Without linking, the annotation simply represents a hypothesis that a given entity can be classed as either an arthropod, a trait, or a value. Taxonomic names are structured terms deriving from academic consensus, they are also relatively consistent and used in a similar format in most publications, therefore normalisation should usually be feasible. Indeed 63% of taxon entities matched terms in the dictionary (Figure 4), *i.e.* they could be linked to species and higher-level taxonomic names from the Catalogue of Life (COL). This result is promising when compared to the performance of other NER systems developed specifically for taxonomic name recognition, which, depending on the corpus used for assessment, can range in precision from 23% to 96% (Le Guillarme and Thuiller 2022). In preparing the arthropod taxa dictionary only COL accepted names were considered (see Methods), however, given the diversity of sources and ages of the processed publications, a universal taxon dictionary including all ever-recorded names and synonyms would have resulted in higher normalisation levels. With respect to traits, normalisation levels were considerably lower than for taxa, reflecting the much less structured manner in which traits are usually described in publications (Figure 4). Of the three categories of traits, morphology achieved the highest level of linking, likely explained by the expectation for taxonomic treatment publications to be dense in morphological descriptions that are used to define and distinguish between species. While nearly a third of the feeding ecology traits from the dictionary could be linked to trait entities in the documents, this represents only a small fraction of all annotated trait entities. In contrast, linking of habitat traits proved very challenging, with only six terms being linked to annotated trait entities. This likely reflects considerable differences between the formal style of term names used by the Environment Ontology and the more variable descriptions of habitats used in natural language. In summary, while entity normalisation remains challenging, iterative extension, refinement, and curation of the dictionaries should lead to higher levels of linking, *e.g.* by expanding the taxon dictionary to include all known names and synonyms, and by extending, revising, and curating the feeding ecology vocabulary and habitat ontology to add synonyms and additional terms that align better with natural language usage. Additionally, while this work focused on arthropod organismal and ecological traits, enhancing the trait dictionaries –which are key for the normalisation process– by aligning them with other existing ontologies developed for biodiversity research more generally, such as the BiodivOnto (Abdelmageed et al. 2021), could improve both entity recognition and the linking of entities to formalised concepts.

### Integrating community feedback into future ArTraDB functionalities

In addition to enhancing the breadth and utility of the trait dictionaries and the performance of the workflow, future work will also need to extend the set of gold standard annotations to improve NER and RE performance through better training and benchmarking. An important source of new annotations could be achieved through further developing the ArTraDB resource functionalities to allow user curation of the workflow-predicted annotations. This would require integration of curation tools into the Annotation Viewer window (Figure 8) that allow community users to edit, add, confirm, or delete entity and relationship annotations. This would facilitate “community curators”, *i.e.*, scientifically literate individuals but not necessarily domain experts, to confirm or modify the annotations, as well as deleting wrong or adding missing annotations, in a crowdsourcing-like operation. The development of this functionality opens up opportunities to then incrementally fine-tune the prediction models by providing them with human-validated annotations as additional training data. While technically feasible and indeed prototyped in the development version of ArTraDB, before deploying such an interactive functionality it would be important to first build the infrastructure for the long-term preservation of community-sourced curated annotations. This could involve the publication of community-confirmed entities and relationships to infrastructures such as the Encyclopedia of Life or Wikidata, however this is technically challenging and would only cover normalised entities. Alternatives could be to publish Journal Article Tag Suite (JATS) XML and/or BioC JSON files on repositories like Zenodo (CERN and OpenAIRE 2024), or even to archive individual annotations as nanopublications with detailed provenance information to identify the source document and location within the document. A more comprehensive approach might instead be to focus on infrastructures supporting the central indexing of biodiversity-related literature (Pasche et al. 2023a, 2023b) that facilitate the addition of annotations to the articles using JATS XML and/or BioC JSON formats. Once a sustainable solution is in place, then ArTraDB could begin to collect community feedback for use as part of improved training and benchmarking datasets, and for collation into versioned annotation sets for archiving or integration into open biodiversity literature services.

### Perspectives on literature mining for biology, ecology, and evolution research

Our methodological explorations to develop tools and resources to advance the use of text mining approaches in biology, ecology, and evolution research, demonstrate the feasibility of semi-automating the building of open databases of organismal and ecological traits extracted from the literature. Even if the annotated arthropod taxon-trait-value triples are sparse, they enable researchers to quickly locate documents pertaining to specific species and traits. This not only accelerates the initial stages of data curation but also points researchers to the exact locations within documents where relevant data can be found, thereby having the potential to enhance the efficiency of research workflows. While there remain several technical challenges to overcome, including how to best leverage the power of modern LLMs in these processes (Farrell et al. 2024, Marcos et al. 2024, Keck et al. 2025), the results provide a framework that could be extended beyond the focus on arthropods. These and other TDM and NLP initiatives in the biodiversity domain will enhance data synthesis studies, make literature reviews more reproducible, greatly facilitate identification of research knowledge gaps and biases, as well as drive data-informed investigations of ecological and evolutionary trends and patterns (Farrell et al. 2022, 2024). When a trait or set of traits has been carefully curated, and the relevant group of species is well-represented with genomics data, researchers can begin to ask how genetic and genomic changes relate to observable phenotypic differences, *e.g.* swallowtail butterfly lineages with host-plant shifts have more genes under positive selection than non-shifting lineages (Allio et al. 2021), and transitions to parthenogenesis (asexual reproduction) in stick insects are accompanied by greatly reduced genetic diversity and reduced rates of positive selection (Jaron et al. 2022). As genomics and other “omics” data become more accessible, and as catalogues of species traits become more comprehensive, –relying on automation for scale and curation for quality control– new opportunities for studying complex evolutionary processes will emerge (Cornwallis and Griffin 2024). There are also implications beyond fundamental research, *e.g.* in the context of detecting and reporting biodiversity change globally, data and knowledge are critical for the measurement framework of Essential Biodiversity Variables (EBVs), where TDM and NLP tools and services could contribute especially to informing “species traits” EBVs (Kissling et al. 2018). While much of the knowledge about biodiversity collected and published over centuries remains largely not machine-readable, digitisation efforts and open science initiatives are contributing to the opening up of biodiversity literature (Agosti et al. 2024). Therefore, the continued development and enhancement of specialist and generalist biodiversity literature mining tools and resources is required to serve researcher needs as well as to inform assessments and guide policy decisions on the protection and restoration of biological diversity for a sustainable future.

## Acknowledgements

The authors thank Morgane Massy from the University of Lausanne for participating in the annotation of the gold-standard set of curated articles used for training and assessing performance, and Guido Sautter from Plazi for preparing and sharing the arthropod taxonomic treatments from TreatmentBank.

## Funding

This work was primarily funded through Swiss National Science Foundation (SNSF) Spark grant 196125 to RMW. The authors acknowledge additional support from SNSF grants 202669 (to RMW).

## Conflict of interest

The authors declare no conflicts of interest.

## Supplementary materials

**Supplementary File S1**: Curated trait dictionaries

An MS Excel spreadsheet presenting the lists of trait dictionaries for feeding ecology, habitat, and morphology, with links to the source resources, synonyms, and definitions.

**Supplementary File S2**: Annotator guidelines

A PDF file of the notes and guidelines developed by the annotators during the curation of the gold-standard annotation data.

**Supplementary File S3**: Gold-standard annotated documents

An MS Excel spreadsheet listing the annotated files, the number of annotations for each type, and the corresponding annotators.

**Supplementary File S4**: NER and RE baseline results

An MS Excel spreadsheet containing five tables with exact scores for all configurations, in terms of recall, precision, and F-score values, along with the corresponding support for each class and the macro and weighted averages.

## Notes

### Competing Interest Statement

The authors have declared no competing interest.

## References

Abdelmageed N, Algergawy A, Samuel S, König-Ries B (2021) BiodivOnto: Towards a core ontology for biodiversity. In: Verborgh R, Dimou A, Hogan A, d’Amato C, Tiddi I, Bröring A, Mayer S, Ongenae F, Tommasini R, Alam M (Eds), The Semantic Web: ESWC 2021 Satellite Events. Lecture Notes in Computer Science. Springer International Publishing, Cham, 3–8. 10.1007/978-3-030-80418-3_1

Abdelmageed N, Löffler F, Feddoul L, Algergawy A, Samuel S, Gaikwad J, Kazem A, König-Ries B (2022) BiodivNERE: Gold standard corpora for named entity recognition and relation extraction in the biodiversity domain. Biodiversity Data Journal 10: e89481. 10.3897/BDJ.10.e89481

Agosti D, Bénichou L, Casino A, Nielsen L, Ruch P, Kishor P, Penev L, Mergen P, Arvanitidis C (2024) Liberate the power of biodiversity literature as FAIR digital objects. Research Ideas and Outcomes 10: e126586. 10.3897/rio.10.e126586

Agosti D, Benichou L, Addink W, Arvanitidis C, Catapano T, Cochrane G, Dillen M, Döring M, Georgiev T, Gérard I, Groom Q, Kishor P, Kroh A, Kvaček J, Mergen P, Mietchen D, Pauperio J, Sautter G, Penev L (2022) Recommendations for use of annotations and persistent identifiers in taxonomy and biodiversity publishing. Research Ideas and Outcomes 8: e97374. 10.3897/rio.8.e97374

Allio R, Nabholz B, Wanke S, Chomicki G, Pérez-Escobar OA, Cotton AM, Clamens A-L, Kergoat GJ, Sperling FAH, Condamine FL (2021) Genome-wide macroevolutionary signatures of key innovations in butterflies colonizing new host plants. Nature Communications 12: 354. 10.1038/s41467-020-20507-3

Bánki O, Roskov Y, Döring M, Ower G, Robles DRH, Corredor CAP, Jeppesen TS, Örn A, Pape T, Hobern D, Garnett S, Little H, DeWalt RE, Ma K, Miller J, Orrell T, Aalbu R, Abbott J, Aedo C, Aescht E, Alexander S, Alonso-Zarazaga MA, Alvarez B, Andrella GC, Antonietto LS, Arango C, Artois T, Burgos MA, Atkinson S, Atwood JJ, Sartori ÂLB, Bailly N, Baixeras J, Baker E, Balan A, Bamber R, Bandyopadhyay S, Barber-James H, Pinto RB, Barrett R, Bartolozzi L, Bartsch I, Beccaloni G, Bellamy CL, Bellan-Santini D, Bellinger PF, Ben-Dov Y, Blasco-Costa I, Boatwright JS, Bock P, Bolton B, Borges LM, Bortoluzzi R, Bossard RL, Bota-Sierra C, Bouchard P, Bourgoin T, Boury-Esnault N, Boxshall G, Boyko C, Brandão S, Braun H, Bray R, Brehm G, Brinda JC, Brock PD, Broich SL, Brown J, Brown S, Bruce N, Brullo S, Bruneau A, Bush L, Büscher T, Błażewicz-Paszkowycz M, Cabras A, Cairns S, Calonje M, Cardinal-McTeague W, Cardoso D, Cardoso L, Castilho RC, Silva ICC, Cervantes A, Chernyshev A, Chevillotte H, Choo LM, Christiansen KA, Cianferoni F, Cigliano MM, Clarke R, Monteiro TC e, Collins A, Compton J, Copilac:-Ciocianu D, Corbari L, Cordeiro R, Cortés-Hernández K, Costello M, Crameri S, Cruz-López JA, Cárdenas P, Daly M, Daneliya M, Dauvin J-C, Davie P, Broyer CD, Grave SD, Lima HCD, Prins JD, Prins WD, Sousa FD, Estrella MD la, DeSalle R, Decker P, Decock W, Delgado-Salinas A, Deliry C, Dellapé PM, Heyer JD, Dijkstra K-D, Dmitriev DA, Dohrmann M, Dorado Ó, Dorkeld F, Downey R, Duan L, Díaz M-C, Eades DC, Egan AN, Eitel M, Nagar AE, Emig CC, Engel MS, Garrote PE, Evans GA, Evenhuis NL, Falcão M, Farruggia F, Fauchald K, Fautin D, Favret C, Fisher B, Fišer C, Forró L, Fortuna-Perez AP, Fortune-Hopkins H, Fritsch P, Froese R, Fuchs A, Fujimoto S, Furuya H, Gagnon E, Garic R, Gasca R, Gattolliat J-L, Gerken S, Lima AG de, Gibson D, Gielis C, Gilligan T, Giribet G, Duque JCG, Gittenberger A, Galdo GG del, Gofas S, Goncharov M, Gondim AI, Goodwin C, Govaerts R, Grabowski M, Granado A de A, Gregório B de S, Grehan JR, Grether R, Grimaldi DA, Gross O, Guerra-García JM, Guglielmone A, Guilbert E, Frøslev TG, Gusenleitner J, Haas F, Hadfield KA, Hajdu E, Hassler M, Hastriter MW, Hauser C, Hausmann A, Hayward BW, Hendrycks E, Henry TJ, Hernandes FA, Hernández-Crespo JC, Hine A, Ho B-C, Hodson A, Hoeksema B, Hoenemann M, Holstein J, Hooge M, Hooper J, Hopkins H, Horak I, Horton T, Hošek J, Hughes C, Hughes L, Huys R, Häuser C, Janssens F, Jaume D, Javadi F, Jazdzewski K, Jersabek CD, Johnson KP, Jordão L, Jóźwiak P, Kajihara H, Kakui K, Kallies A, Kamiński MJ, Kanda K, Karanovic I, Kathirithamby J, Kelly M, Kim Y-H, King R, Kirk P, Kitching I, Klautau M, Klitgaard BB, Koenemann S, Korovchinsky NM, Kotov A, Kramina T, Krapp-Schickel T, Kremenetskaia A, Krishna K, Krishna V, Kroh A, Kroupa AS, Kury AB, Kury MS, Kvaček J, Lachenaud O, Lado C, Lambert G, Atunes LLC, Lavin M, Lazarus D, Coze FL, Roux ML, LeCroy S, Linares JL, Lee S, Leitner MF, Lewis GP, Li S-J, Li-Qiang J, Lichtwardt R(†), Lim S-C, Littlewood T, Lohrmann V, Longhorn SJ, Lorenz W, Lowry J, Lozano F, Lumen R, Lyal CH, Lörz A-N, Madin L, Magnien P, Mah C, Mal N, Mamos T, Manconi R, Mansano V, Markello K, Martens K, Martin JH, Martin P, Mashego KS, Maslakova S, Maslin B, Mattapha S, McFadden C, McKamey S, McMurtry JA, Medrano MA, Mees J, Mendes AC, Merrin K, Mesa NC, Messing C, Mielke CGC, Migeon A, Miller DR, Mills C, Minelli A, Mitchell D, Molodtsova T, Valls JFM, Mooi R, Morandini A, Rocha RM da, Morrow C, Moteetee A, Murillo-Ramos L, Murphy B, Narita JPZ, Nery DG, Neu-Becker U, Neuhaus B, Newton A, Lin PNK, Nicolson D, Nielsen JE, Nijhof A, Nishikawa T, Norenburg J, O’Hara T, Ochoa R, Ohashi H, Ohashi K, Ollerenshaw J, Oosterbroek P, Opresko D, Osborne R, Osigus H-J, Oswald JD, Ota Y, Otte D, Ouvrard D, Queiroz LP de, Pandey A, Paulay G, Paulson D, Pauly D, Pennington RT, Pereira J da S, Perez-Gelabert D, Petrusek A, Phillipson P, Pinheiro U, Morim MP, Pisera A, Pitkin B, Plotkin D, Pierezan BP, Poore G, Povydysh M, Praxedes RA, Pulawski WJ, Pyle R, Pühringer F, Rajaei H, Rakotonirina N, Ramos G, Rando J, Filardi FR, Raz L, Read G, Rees T, Reich M, Reimer JD, Rein JO, Reynolds J, Rincón J, Rius M, Robertson T, Robinson G, Robinson GS(†), Rodríguez E, Ruggiero M, Ríos P, Rützler K, Sanborn A, Sanjappa M, Santos SG, Santos-Guerra A, Sartori M, Sattler K, Schierwater B, Schilling S, Schley R, Schmid-Egger C, Schmidt-Rhaesa A, Schoolmeesters P, Schorr M, Schrire B, Schuchert P, Schuh RT, Schönberg C, Rodrigues RS, Scoble M, Seijo G, Seleme EP, Senna A, Serejo C, Sforzi A, Shenkar N, Shimizu G, Siegel V, Sierwald P, Sihvonen P, Flores AS, Carvalho CS de, Simon MF, Simonsen T, Simpson CE, Sinniger F, Sirichamorn Y, Skvarla M, Smith AD, Smith VS, Gissi DS, Sokoloff D, Sotuyo S, Soulier-Perkins A, South EJ, Souza-Filho JF, Spearman L, Spelda J, Steiner A, Stemme T, Sterrer W, Stevenson D, Stiewe MBD, Stirton CH, Straub S, Stueber G, Stöhr S, Subramaniam S, Swalla B, Swedo J, Sánchez-Ruiz M, Sørensen MV, Taiti S, Takiya DM, Tandberg AH, Tavakilian G, Taylor K, Thessen A, Thomas JD, Thomas P, Thomson S, Thuesen E, Thulin M, Thurston M, Thuy B, Todaro A, Torke BM, Tsai S-Y, Turiault M, Turner JRG, Turner T, Turon X, Tyler S, Uetz P, Ulmer JM, Vacelet J, Vachard D, Vader W, Domedel GV, Burgt XV der, Vandepitte L, Vanhoorne B, Vatanparast M, Verhoeff T, Vonk R, Väinölä R, Walker-Smith G, Walter TC, Wambiji N, Wanke D, Watling L, Weaver H, Webb J, Welbourn WC, Whipps C, White K, Wilding N, Williams G, Wilson AJG, Wing P, Winitsky S, Wirth CC, Wojciechowski M, Woodman S, Xavier J, Yi T, Yoder M, Yu DSK, Yunakov N, Zahniser J, Zeidler W, Zhang R, Zhang ZQ, Zinetti F, d’Hondt J-L, Moraes GJ de, Oliveira ABR de, Voogd N de, Río MG del, Haaren T van, Nieukerken EJ van, Ofwegen L van, Soest R van, Şentürk O (2024) Catalogue of Life. Version 2024-12-19. 10.48580/dglq4

Basaldella M, Furrer L, Tasso C, Rinaldi F (2017) Entity recognition in the biomedical domain using a hybrid approach. Journal of Biomedical Semantics 8: 51. 10.1186/s13326-017-0157-6

Buttigieg PL, Pafilis E, Lewis SE, Schildhauer MP, Walls RL, Mungall CJ (2016) The environment ontology in 2016: bridging domains with increased scope, semantic density, and interoperation. Journal of Biomedical Semantics 7: 57. 10.1186/s13326-016-0097-6

Cejuela JM, McQuilton P, Ponting L, Marygold SJ, Stefancsik R, Millburn GH, Rost B, the FlyBase Consortium (2014) Tagtog: Interactive and text-mining-assisted annotation of gene mentions in PLOS full-text articles. Database 2014: bau033–bau033. 10.1093/database/bau033

CERN, OpenAIRE (2024) Zenodo. 10.25495/7GXK-RD71

Chang A, Jeske L, Ulbrich S, Hofmann J, Koblitz J, Schomburg I, Neumann-Schaal M, Jahn D, Schomburg D (2021) BRENDA, the ELIXIR core data resource in 2021: new developments and updates. Nucleic Acids Research 49: D498–D508. 10.1093/nar/gkaa1025

Church SH, Donoughe S, De Medeiros BAS, Extavour CG (2019) A dataset of egg size and shape from more than 6,700 insect species. Scientific Data 6: 104. 10.1038/s41597-019-0049-y

Cohen J (1960) A coefficient of agreement for nominal scales. Educational and Psychological Measurement 20: 37–46. 10.1177/001316446002000104

Comeau DC, Islamaj Dogan R, Ciccarese P, Cohen KB, Krallinger M, Leitner F, Lu Z, Peng Y, Rinaldi F, Torii M, Valencia A, Verspoor K, Wiegers TC, Wu CH, Wilbur WJ (2013) BioC: a minimalist approach to interoperability for biomedical text processing. Database 2013: bat064–bat064. 10.1093/database/bat064

Cornwallis CK, Griffin AS (2024) A guided tour of phylogenetic comparative methods for studying trait evolution. Annual Review of Ecology, Evolution, and Systematics 55: 181–204. 10.1146/annurev-ecolsys-102221-050754

Devlin J, Chang M-W, Lee K, Toutanova K (2019) BERT: Pre-training of deep bidirectional transformers for language understanding. Proceedings of the 2019 Conference of the North: 4171–4186. 10.18653/v1/N19-1423

Farrell MJ, Brierley L, Willoughby A, Yates A, Mideo N (2022) Past and future uses of text mining in ecology and evolution. Proceedings of the Royal Society B: Biological Sciences 289: 20212721. 10.1098/rspb.2021.2721

Farrell MJ, Le Guillarme N, Brierley L, Hunter B, Scheepens D, Willoughby A, Yates A, Mideo N (2024) The changing landscape of text mining: a review of approaches for ecology and evolution. Proceedings of the Royal Society B: Biological Sciences 291: 20240423. 10.1098/rspb.2024.0423

Feron R, Waterhouse RM (2022a) Assessing species coverage and assembly quality of rapidly accumulating sequenced genomes. GigaScience 11: giac006. 10.1093/gigascience/giac006

Feron R, Waterhouse RM (2022b) Exploring new genomic territories with emerging model insects. Current Opinion in Insect Science 51: 100902. 10.1016/j.cois.2022.100902

Furrer L, Cornelius J, Rinaldi F (2022) Parallel sequence tagging for concept recognition. BMC Bioinformatics 22: 623. 10.1186/s12859-021-04511-y

Gaimari SD (2017) The dipteran family Celyphidae in the New World, with discussion of and key to world genera (Insecta, Diptera). ZooKeys 711: 113–130. 10.3897/zookeys.711.20840

Grimaldi DA, Engel MS (2005) Evolution of the insects. Cambridge university press, Cambridge.

Guidoti M, Sokolowicz C, Simoes F, Gonçalves V, Ruschel T, Alvares D, Agosti D (2021) TreatmentBank: Plazi’s strategies and its implementation to most efficiently liberate data from scholarly publications. Biodiversity Information Science and Standards 5: e75690. 10.3897/biss.5.75690

Hedrick BP, Heberling JM, Meineke EK, Turner KG, Grassa CJ, Park DS, Kennedy J, Clarke JA, Cook JA, Blackburn DC, Edwards SV, Davis CC (2020) Digitization and the future of natural history collections. BioScience 70: 243–251. 10.1093/biosci/biz163

Jaron KS, Parker DJ, Anselmetti Y, Tran Van P, Bast J, Dumas Z, Figuet E, François CM, Hayward K, Rossier V, Simion P, Robinson-Rechavi M, Galtier N, Schwander T (2022) Convergent consequences of parthenogenesis on stick insect genomes. Science Advances 8: eabg3842. 10.1126/sciadv.abg3842

Keck F, Broadbent H, Altermatt F (2025) Extracting massive ecological data on state and interactions of species using large language models. 10.1101/2025.01.24.634685

Kim Sang EFT, De Meulder F (2003) Introduction to the CoNLL-2003 shared task: Language-independent named entity recognition. In: Proceedings of the Seventh Conference on Natural Language Learning at HLT-NAACL 2003. , 142–147. Available from: https://aclanthology.org/W03-0419/.

Kissling WD, Walls R, Bowser A, Jones MO, Kattge J, Agosti D, Amengual J, Basset A, Van Bodegom PM, Cornelissen JHC, Denny EG, Deudero S, Egloff W, Elmendorf SC, Alonso García E, Jones KD, Jones OR, Lavorel S, Lear D, Navarro LM, Pawar S, Pirzl R, Rüger N, Sal S, Salguero-Gómez R, Schigel D, Schulz K-S, Skidmore A, Guralnick RP (2018) Towards global data products of Essential Biodiversity Variables on species traits. Nature Ecology & Evolution 2: 1531–1540. 10.1038/s41559-018-0667-3

Le Guillarme N, Thuiller W (2022) TaxoNERD: Deep neural models for the recognition of taxonomic entities in the ecological and evolutionary literature. Methods in Ecology and Evolution 13: 625–641. 10.1111/2041-210X.13778

Lee J, Yoon W, Kim S, Kim D, Kim S, So CH, Kang J (2020) BioBERT: a pre-trained biomedical language representation model for biomedical text mining. Wren J (Ed.). Bioinformatics 36: 1234–1240. 10.1093/bioinformatics/btz682

Lever J, Altman R, Kim J-D (2020) Extending TextAE for annotation of non-contiguous entities. Genomics & Informatics 18: e15. 10.5808/GI.2020.18.2.e15

Li Y, Ramprasad R, Zhang C (2024) A simple but effective approach to improve structured language model output for information extraction. 10.48550/ARXIV.2402.13364

Liu Y, Ott M, Goyal N, Du J, Joshi M, Chen D, Levy O, Lewis M, Zettlemoyer L, Stoyanov V (2019) RoBERTa: A robustly optimized BERT pretraining approach. Available from: http://arxiv.org/abs/1907.11692 (December 16, 2024).

Mammola S, Pavlek M, Huber BA, Isaia M, Ballarin F, Tolve M, Čupić I, Hesselberg T, Lunghi E, Mouron S, Graco-Roza C, Cardoso P (2022) A trait database and updated checklist for European subterranean spiders. Scientific Data 9: 236. 10.1038/s41597-022-01316-3

Marcos D, van de Vlasakker R, Athanasiadis IN, Bonnet P, Goeau H, Joly A, Kissling WD, Leblanc C, van Proosdij ASJ, Panousis KP (2024) Fully automatic extraction of morphological traits from the Web: utopia or reality? 10.48550/ARXIV.2409.17179

McCallen E, Knott J, Nunez-Mir G, Taylor B, Jo I, Fei S (2019) Trends in ecology: shifts in ecological research themes over the past four decades. Frontiers in Ecology and the Environment 17: 109–116. 10.1002/fee.1993

McHugh ML (2012) Interrater reliability: the kappa statistic. Biochemia Medica 22: 276–282.

Montagna M (2011) Pachybrachis sassii, a new species from the Mediterranean Giglio Island (Italy) (Coleoptera, Chrysomelidae, Cryptocephalinae). ZooKeys 155: 51–60. 10.3897/zookeys.155.1951

Montani I, Honnibal M, Boyd A, Van Landeghem S, Peters H (2023) explosion/spaCy: v3.7.2: Fixes for APIs and requirements. 10.5281/ZENODO.1212303

Mündler N (2024) nielstron/quantulum3. Available from: https://github.com/nielstron/quantulum3 (November 25, 2024).

Mungall C, Matentzoglu N, Balhoff J, Osumi-Sutherland D, Duncan B, Pgaudet, Tan S, Hoyt CT, Pilgrim C, Overton JA, Lauren, Caron A, Nomi Harris, Moxon S, Lschriml, Vasilevsky N, Toro S, Goutte-Gattat D, Brush M, Vasundra Touré, Bretaudeau A, Cain S, Haendel M, DiatomsRcool, Bide Zhang, Dowland C, Dooley D, Actions-User, Hammock J (2023) The OBO relation ontology, http://purl.obolibrary.org/obo/ro.owl. 10.5281/ZENODO.593101

Mungall CJ, Torniai C, Gkoutos GV, Lewis SE, Haendel MA (2012) Uberon, an integrative multi-species anatomy ontology. Genome Biology 13: R5. 10.1186/gb-2012-13-1-r5

Parr CS, Wilson N, Leary P, Schulz K, Lans K, Walley L, Hammock J, Goddard A, Rice J, Studer M, Holmes J, Corrigan, Jr. R (2014) The Encyclopedia of Life v2: Providing global access to knowledge about life on Earth. Biodiversity Data Journal 2: e1079. 10.3897/BDJ.2.e1079

Pasche E, Agosti D, Penev L, Groom Q, Flament A, Gobeill J, Ruch P (2023a) Towards “Biodiversity PMC.” Biodiversity Information Science and Standards 7: e111647. 10.3897/biss.7.111647

Pasche E, Gobeill J, Agosti D, Penev L, Groom Q, Georgiev T, Gaillac E, Flament A, Caucheteur D, Michel P-A, Ruch P (2023b) From SIBiLS to Biodiversity PMC: Foundations for the One Health Library. Biodiversity Information Science and Standards 7: e111660. 10.3897/biss.7.111660

Ramshaw LA, Marcus MP (1999) Text chunking using transformation-based learning. In: Armstrong S, Church K, Isabelle P, Manzi S, Tzoukermann E, Yarowsky D (Eds), Natural Language Processing Using Very Large Corpora. Text, Speech and Language Technology. Springer Netherlands, Dordrecht, 157–176. 10.1007/978-94-017-2390-9_10

Rosonovski S, Levchenko M, Ide-Smith M, Faulk L, Harrison M, McEntyre J (2023) Searching and evaluating publications and preprints using Europe PMC. Current Protocols 3: e694. 10.1002/cpz1.694

Shirey V, Larsen E, Doherty A, Kim CA, Al-Sulaiman FT, Hinolan JD, Itliong MGA, Naive MAK, Ku M, Belitz M, Jeschke G, Barve V, Lamas G, Kawahara AY, Guralnick R, Pierce NE, Lohman DJ, Ries L (2022) LepTraits 1.0 A globally comprehensive dataset of butterfly traits. Scientific Data 9: 382. 10.1038/s41597-022-01473-5

Stork NE (2018) How many species of insects and other terrestrial arthropods are there on Earth? Annual Review of Entomology 63: 31–45. 10.1146/annurev-ento-020117-043348

Wong MKL, Guénard B, Lewis OT (2019) Traitc:based ecology of terrestrial arthropods. Biological Reviews 94: 999–1022. 10.1111/brv.12488

Yamada I, Asai A, Shindo H, Takeda H, Matsumoto Y (2020) LUKE: Deep contextualized entity representations with entity-aware self-attention. In: Proceedings of the 2020 Conference on Empirical Methods in Natural Language Processing (EMNLP). Association for Computational Linguistics, Online, 6442–6454. 10.18653/v1/2020.emnlp-main.523

