## Supplementary File S2 for "From literature to biodiversity data: mining arthropod organismal and ecological traits with machine learning"

### ArTraDB notes on the process of annotating the documents, and descriptions of guidelines for dealing with some of the more complex cases

To fine-tune and formally evaluate the performance of the NLP models employed for the Named Entity Recognition and Relationship Extraction tasks, two entomology domain experts annotated a set of 25 articles randomly selected from the 3’650 obtained from PMC. The two annotators employed the tagtog text annotation tool (Cejuela et al. 2014) that provides a user-friendly interface to manually annotate and normalise entities in documents imported from PMC, as well as to add entity labels, relationships, and more. The annotators worked independently on their assigned documents to avoid biasing each other, however, they did develop a set of guidelines during the annotation process to describe the steps to follow for dealing with some more complex cases.

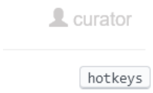


Check out the hotkeys box for shortcuts: top-right of page
t = add relation
d = delete annotation

#### In no particular order:

- **[1] How to deal with places/regions?**

Technically it seems like it should be:
Arthropod <=> has_trait [habitat/location] <=> has_value [the place/region]
But often the ‘habitat/location’ trait is very hard to define, e.g. ‘from’ ‘found in’ etc.
So instead we go directly and put the place/region as the TRAIT with a relation to the ARTHROPOD rather than a VALUE with a relation to a ‘habitat/location’ TRAIT

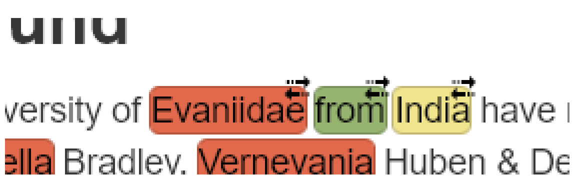


**ARTHROPOD ⇔ linker: ‘from’ ‘distribution’ ‘located’ ‘inhabits’ ‘found in’ ‘lives in’ ⇔ VALUE (e.g. geographic location - forest, river, etc.)**

NB: Because ‘linker’ terms can be very generic (e.g. from) best to turn off pre-selections feature before annotating them

- **[2] How to deal with adjectives of traits?**

E.g. metathoracic femur - the femur of the metathorax
In such cases the qualifier clearly adds precision so we should try to capture these cases wherever possible, i.e. 'metathoracic femur' is the TRAIT, not just 'femur'
Or should we annotate both?
=> no, always go for the most specific TRAIT possible

- **[3] When only pairs are possible - what to do?**

Often we can only make a single pairwise relation between ARTHROPOD and TRAIT
I.e. a VALUE that would complete the triple ARTHROPOD-TRAIT-VALUE is hard to annotate
Knowing that the TRAIT is mentioned in the document and can be linked to an ARTHROPOD is still valuable, even if we cannot annotate any values associated with the pair, therefore still try to annotate valid ARTHROPOD ⇔ TRAIT pairs

- **[4] How to deal with qualifiers of taxa?**

A good example is when when sex or life stage is mentioned
- do we annotate BODY LENGTH or FEMALE BODY LENGTH?
=> no, we add a QUALIFIER to the TAXON
E.g. Here we have to link female to Zeuxevania hubeni using a QUALIFIER (BLUE) relation

**New information**

A new evaniid species Zeuxevania hubeni sp. nov. is described, based on a female specimen collected from Kadaludi Bird Sanctuary, Kerala, India. The new species is compared with Z. curvicarinata (Cameron), as well as Z. kasauliensis (Muzaffer) and a key to Indian species, based on females, is provided. The type specimen is deposited in the National Zoological Collection, Zoological Survey of India, Kolkata, India.

- **[5] When no relations are possible - what to do?**

Often we can annotate ARTHROPOD, or TRAIT, and occasionally VALUE, but adding relations is not possible
ARTHROPOD => yes, useful to know which arthropods are mentioned in the document
TRAIT => yes, useful to know the document mentions the trait, even if we cannot connect it to a specific arthropod
VALUE => no, values are usually too generic and thus on their own are not useful
Example:

**Materials and methods**

The specimen was collected using a sweep net from Kadaludi Bird Sanctuary, Kerala, India, killed by ethyl acetate and stored in 70% ethyl alcohol. The specimen was later dried and mounted on a rectangular card using water-soluble glue. Photographs were taken with a Nikon DS-Ri2 camera, mounted on a Nikon SMZ25 stereozoom microscope and processed by the NIS-Elements BR Analysis v5.20.00 software. Generic placement was determined using the key provided by Deans and Huben (2003). The examined holotype is deposited in the National Zoological Collection, Zoological Survey of India, Kolkata, India (NZC). The following abbreviations are used in the text: OOL – Minimum distance between the posterior ocellus and eye margin; POL – Minimum distance between the two posterior ocelli; OAL – Minimum distance between the posterior ocellus and anterior ocellus; F1-F11 – Funicular segments 1-11.

- **[6] Another example of a potential QUALIFIER**

Here the 'exserted part' refers to a quality of the ovipositor
⇒ Just annotate this as a QUALIFIER

; ovipositor (exserted part) length, 0.35.

- **[7] How to deal with relations that could be annotated several times?**

Example: Zeuxevania ⇔ from ⇔ India can be annotated twice in this paragraph
Is it still useful to annotate the same relations again?
⇒ YES, always annotate all occurrences locally

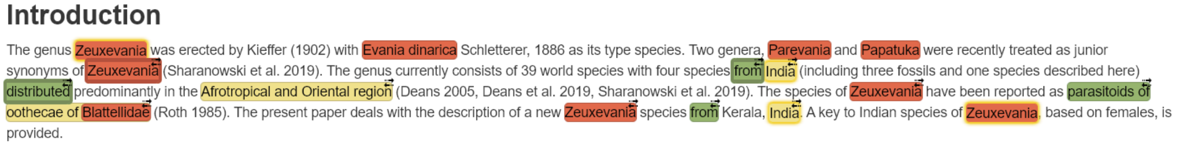
**Introduction**

The genus Zeuxevania was erected by Kieffer (1902) with Evania dinarica Schletterer, 1886 as its type species. Two genera, Parevania and Papatuka were recently treated as junior synonyms of Zeuxevania (Sharanowski et al. 2019). The genus currently consists of 39 world species with four species from India (including three fossils and one species described here) distributed predominantly in the Afrotropical and Oriental region (Deans 2005, Deans et al. 2019, Sharanowski et al. 2019). The species of Zeuxevania have been reported as parasitoids of oothecae of Blattellidae (Roth 1985). The present paper deals with the description of a new Zeuxevania species from Kerala, India. A key to Indian species of Zeuxevania, based on females, is provided.

- **[8] When an arthropod is a value of a trait?**

The species of Zeuxevania have been reported as parasitoids of oothecae of Blattellidae
Here Blattellidae is both an arthropod (group) and a value
Zeuxevania <=> parasitoids of <=> oothecae  NO: use the full phrase as VALUE
Zeuxevania <=> parasitoids of <=> Blattellidae  NO: use the full phrase as VALUE

**Zeuxevania <=> parasitoids of <=> oothecae of Blattellidae**

Also annotate the ARTHROPOD

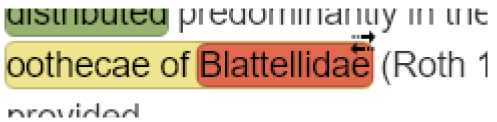


VALUE = oothecae of Blattellidae (yellow)
ARTHROPOD = Blattellidae (red, here shown on top of yellow)

- **[9] When more than one adjective is linked to a trait?**

E.g. Fore and mid coxa, orange brown

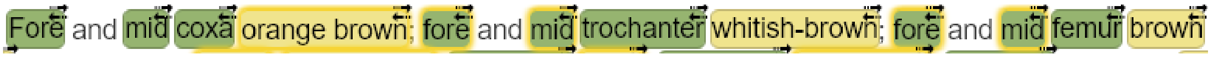


As above? YES

- Annotate 3 TRAITs Fore, mid, & coxa
- Link coxa to ARTHROPOD and to TRAIT (orange brown)
- Link coxa to mid
- Link coxa to Fore

Or a related case, two body parts with one adjective?

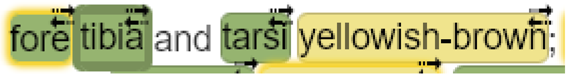


fore ⇔ tibia
fore ⇔ tarsi
tibia ⇔ yellowish-brown
tarsi ⇔ yellowish-brown

- **[10] When Genus and species both appear but not both annotated?**

E.g. here Evania was annotated because it appeared earlier in the document, so now we should delete the annotation on the Genus and make a new annotation spanning both Genus and species

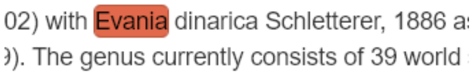

=====>
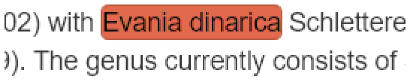


- **[11] When trait words are split and/or composite?**

Example: head width (height), 0.9 (0.1)

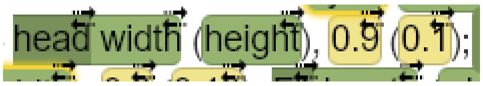


TRAITS

- head
- head width
- height

VALUES

- 0.9
- 0.1

RELATIONS

- head width ⇔ 0.9
- head ⇔ height (TRAIT-TRAIT relation)
- head ⇔ 0.1

Example: eye length (width), 0.6 (0.3)

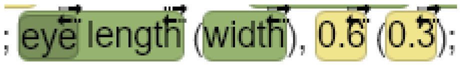


TRAITS

- eye
- eye length
- width

VALUES

- 0.6
- 0.3

RELATIONS

- eye length ⇔ 0.6
- eye ⇔ width (TRAIT-TRAIT relation)
- eye ⇔ 0.3

NOTE: Discontinuous trait-trait relations should not contain more than 2 entities

- **[12] Traits with 2 or more measurements**

Here we have length (width) for several traits in a row

Option 1:
Arthropod ⇔ scape  TAXON-TRAIT
scape ⇔ 163   TRAIT to VALUE-1
scape ⇔ 50    TRAIT to VALUE-2
scape ⇔ length   TRAIT-TRAIT-1
scape ⇔ width    TRAIT-TRAIT-2
No way of telling which value corresponds to length and which to width

; F3 with dorsal margin 3.21× as long as wide, and 0.6× as long as clava; clava 3.95× as long as wide, pointed and distinctly curved ventrally at apex. Measurements, length (width): scape, 163 (50); pedicel, 75 (35); F1, 28 (20); F2, 40 (20); F3, 90 (28); clava 150 (38).

Or option 2:
Arthropod ⇔ scape  TAXON-TRAIT
scape ⇔ 163 (50)   TRAIT to VALUE-1+2
scape ⇔ length (width)   TRAIT-TRAIT-1+2

Or option 3 (The order is important. If we link length/width to the value, we can then unambiguously determine the relationship afterwards.):
Arthropod ⇔ scape  TAXON-TRAIT
scape ⇔ length   TRAIT-TRAIT-1
scape ⇔ width    TRAIT-TRAIT-2
length ⇔ 163   TRAIT to VALUE-1
width ⇔ 50    TRAIT to VALUE-2

The problem here - as soon as there is another body part we again have ambiguity
⇒ because length and width will be associated with several values and several body parts
scape ⇔ length ⇔ 163     pedicel ⇔ length ⇔ 75     F1 ⇔ length ⇔ 28
Length will be linked to 163, 75, and 28
Length will be linked to scape, pedicel, and F1

SOLUTION

; F3 with dorsal margin 3.21× as long as wide, and 0.6× as long as clava; clava 3.95× as long as wide, pointed and distinctly curved ventrally at apex. Measurements, length (width): scape, 163 (50); pedicel, 75 (35); F1, 28 (20); F2, 40 (20); F3, 90 (28); clava 150 (38).

Annotate length (width) as a combined trait
Annotate each pair of values as a combined value, e.g. 163 (50)
Then add relations
ARTHROPOD ⇔ scape
ARTHROPOD ⇔ pedicel
ARTHROPOD ⇔ F1
ARTHROPOD ⇔ F2
ARTHROPOD ⇔ F3
ARTHROPOD ⇔ clava
scape ⇔ length (width)

pedicel ⇔ length (width)
F1 ⇔ length (width)
F2 ⇔ length (width)
F3 ⇔ length (width)
clava ⇔ length (width)
scape ⇔ 163 (50)
pedicel ⇔ 75 (35)
F1 ⇔ 28 (20)
F2 ⇔ 40 (20)
F3 ⇔ 90 (28)
clava ⇔ 150 (38)
